# Whole genome doubling confers unique genetic vulnerabilities on tumor cells

**DOI:** 10.1101/2020.06.18.159095

**Authors:** Ryan J. Quinton, Amanda DiDomizio, Marc A. Vittoria, Carlos J. Ticas, Sheena Patel, Yusuke Koga, Kristýna Kotýnková, Jasmine Vakhshoorzadeh, Nicole Hermance, Taruho S. Kuroda, Neha Parulekar, Alison M. Taylor, Amity L. Manning, Joshua D. Campbell, Neil J. Ganem

**Affiliations:** Department of Pharmacology & Experimental Therapeutics, Boston University School of Medicine, Boston, MA, USA; Department of Medicine, Boston University School of Medicine, Boston, MA 02118, USA; Department of Biology and Biotechnology, Worcester Polytechnic Institute, Worcester, MA; Department of Molecular Pathobiology and Cell Adhesion Biology, Mie University Graduate School of Medicine, Mie, Japan; Department of Medical Oncology, Dana-Farber Cancer Institute, Boston, MA, USA; Cancer Program, Broad Institute, Cambridge, MA, USA; Department of Pathology and Cell Biology, Columbia University Medical Center, Member, Herbert Irving Comprehensive Cancer Center, New York, NY, USA

## Abstract

Whole genome doubling (WGD) occurs early in tumorigenesis and generates genetically unstable tetraploid cells that fuel tumor development. Cells that undergo WGD (WGD^+^) must adapt to accommodate their abnormal tetraploid state; however, the nature of these adaptations, and whether they confer vulnerabilities that can subsequently be exploited therapeutically, is unclear. Using sequencing data from ∼10,000 primary human cancer samples and essentiality data from ∼600 cancer cell lines, we show that WGD gives rise to common genetic traits that are accompanied by unique vulnerabilities. We reveal that WGD^+^ cells are more dependent on spindle assembly checkpoint signaling, DNA replication factors, and proteasome function than WGD^−^ cells. We also identify *KIF18A*, which encodes for a mitotic kinesin, as being specifically required for the viability of WGD^+^ cells. While loss of KIF18A is largely dispensable for accurate chromosome segregation during mitosis in WGD^−^ cells, its loss induces dramatic mitotic errors in WGD^+^ cells, ultimately impairing cell viability. Collectively, our results reveal new strategies to specifically target WGD^+^ cancer cells while sparing the normal, non-transformed WGD^−^ cells that comprise human tissue.

---

The vast majority of human cells are diploid and numerous cell cycle controls exist to help ensure that this state is maintained across successive cell divisions^1^. Despite these controls, errors can occur that result in a whole genome doubling (WGD), in which a natively diploid cell transitions to a tetraploid state^1-3^. It has been demonstrated that cells that have experienced a WGD event (hereafter WGD^+^) are oncogenic and can facilitate tumorigenesis^4,5^. WGD promotes tumorigenesis in at least two ways: first, proliferating WGD^+^ cells are genomically unstable and rapidly accumulate both numerical and structural chromosomal abnormalities^5^, and second, WGD^+^ cells are better able to buffer against the negative effects of deleterious mutations and ongoing chromosome instability^6-10^. Such traits enable nascent WGD^+^ tumor cells to proliferate in the presence of otherwise lethal genomic alterations while simultaneously sampling increased genetic permutations, ultimately enabling phenotypic leaps that give rise to tumors^8,11^. WGD also carries important clinical implications, with recent reports showing its correlation with advanced metastatic disease and a worse overall prognosis^12,13^.

Given the oncogenic potential associated with WGD, tumor suppression mechanisms exist to limit the proliferation of these unstable cells. WGD^+^ cells activate both the p53 and Hippo tumor suppressor pathways and are prone to apoptosis, senescence, and immune clearance^14-16^. WGD also gives rise to numerous abnormalities in cellular physiology that impair fitness^6,14,17^. Therefore, in order to promote tumorigenesis, WGD^+^ cells must adapt to overcome these barriers^5,14,18,19^. Thus, while WGD confers traits that favor tumorigenesis, it also imposes adaptive requirements upon cells that could give rise to unique vulnerabilities^20,21^. Identifying and exploiting these vulnerabilities represents an exciting therapeutic avenue, particularly because WGD is broadly shared across multiple tumor types and is a distinguishing characteristic of many tumors^12,22^.

## Identifying genetic alterations enriched in WGD^+^ tumors

To understand the genetic differences between WGD^+^ and WGD^−^ tumors, we first obtained WGD status calls made by the ABSOLUTE algorithm on ∼10,000 primary tumor samples spanning 32 distinct tumor types from The Cancer Genome Atlas (TCGA). This allowed us to separate tumor samples by whether they had (WGD^+^) or had not (WGD^−^) undergone a WGD event^23^. Consistent with previous estimates, we found that ∼36% of tumors experienced at least one WGD during their evolution^12,24^. We also observed a significant range in the occurrence of WGD between different tumor subtypes, implying that specific genetic, physiological, and/or microenvironmental cues can favor or repress WGD-driven tumorigenesis (Fig. 1a).

**Figure 1.**
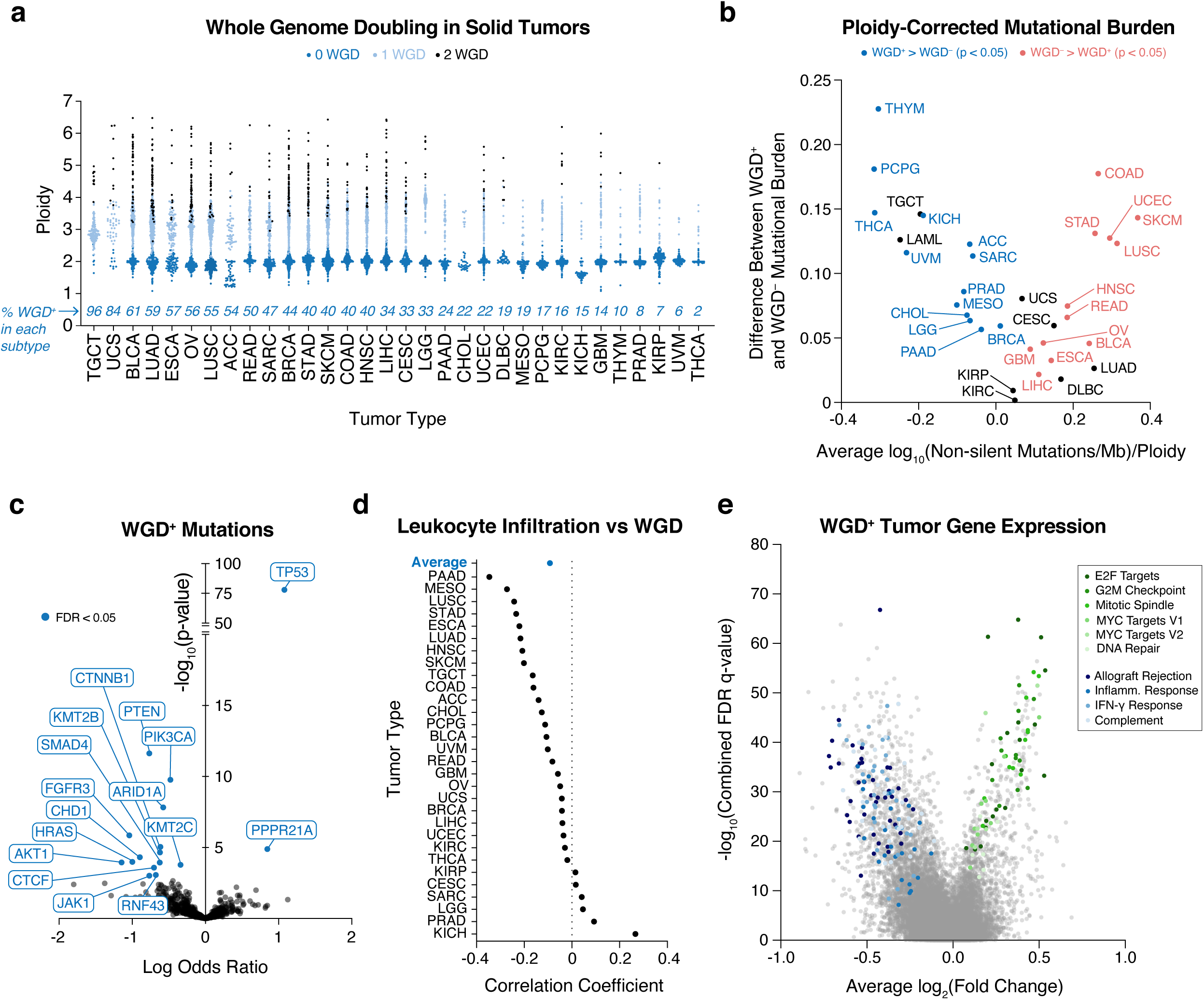
Genetic analysis of WGD^+^ tumors. **(a)** Quantification of WGD status and total ploidy of 9,700 primary human tumor samples from the TCGA using ABSOLUTE. **(b)** Average ploidy-corrected mutational burden in indicated subtypes plotted against the difference in the ploidy-corrected mutational burden between WGD^+^ and WGD^−^ tumors within each subtype (Wilcoxon rank-sum test). **(c)** Enrichment of mutations in WGD^+^ tumors (logistic regression). **(d)** Correlation of leukocyte infiltration and WGD (Pearson’s correlation). **(e)** Gene expression fold changes in WGD^+^ tumors relative to WGD^−^ tumors plotted against combined FDR values across all tumor types with select hits from most significantly enriched gene sets highlighted. * p < 0.05, ** p < 0.01, *** p < 0.001, **** p < 0.0001. TCGA Study Abbreviations: **ACC**-Adrenocortical carcinoma; **BLCA**-Bladder Urothelial Carcinoma; **ESCA**-Esophageal carcinoma; **BRCA**-Breast invasive carcinoma; **CESC**-Cervical squamous cell carcinoma and endocervical adenocarcinoma; **CHOL**-Cholangiocarcinoma; **COAD**-Colon adenocarcinoma; **DLBC**-Lymphoid Neoplasm Diffuse Large B-cell Lymphoma; **GBM**-Glioblastoma multiforme; **HNSC**-Head and Neck squamous cell carcinoma; **KICH**-Kidney Chromophobe; **KIRC**-Kidney renal clear cell carcinoma; **KIRP**-Kidney renal papillary cell carcinoma; **LGG**-Brain Lower Grade Glioma; **LIHC**-Liver hepatocellular carcinoma; **LUAD**-Lung adenocarcinoma; **LUSC**-Lung squamous cell carcinoma; **MESO**-Mesothelioma; **OV**-Ovarian serous cystadenocarcinoma; **PAAD**-Pancreatic adenocarcinoma; **PCPG**-Pheochromocytoma and Paraganglioma; **PRAD**-Prostate adenocarcinoma; **READ**-Rectum adenocarcinoma; **SARC**-Sarcoma; **SKCM**-Skin Cutaneous Melanoma; **STAD**-Stomach adenocarcinoma; **TGCT**-Testicular Germ Cell Tumors; **THYM**-Thymoma; **THCA**-Thyroid carcinoma; **UCS**-Uterine Carcinosarcoma; **UCEC**-Uterine Corpus Endometrial Carcinoma; **UVM**-Uveal Melanoma

Having differentiated WGD^+^ and WGD^−^ tumors, we sought to assess the mutational burden of each cohort in a pan-cancer analysis. We compared the ploidy-corrected mutational burden between WGD^+^ and WGD^−^ tumors and found them to be slightly higher in WGD^+^ tumors (Extended Data Fig. 1a). We also observed that tumors with microsatellite instability (MSI) or mutations in DNA polymerase ε *(POLE)*, which have a very high mutational burden, tend not to experience WGD events, which has been shown in other cohorts^10,12,25,26^. Indeed, only 12/178 tumors we identified as MSI-high/*POLE*-mutated in the TCGA database were WGD^+^ (Extended Data Fig. 1b). Examination within each tumor subtype demonstrated more clearly that WGD^+^ tumors tend to have a higher total mutational burden{Bielski, 2018, Genome doubling shapes the evolution and prognosis of advanced cancers}. However, when we examined the ploidy-corrected mutational burden within each tumor subtype, we found that tissue-specific pressures may differentially affect the acquisition of mutations in WGD^−^ and WGD^+^ tumors (Extended Data Fig. 1c-d). Notably, there were several tumor subtypes where the WGD^−^ tumors had a higher ploidy-corrected mutational burden than the WGD^+^ tumors within that subtype. This tended to occur in subtypes with a high mutational load, characteristic of tumor types prone to MSI or exposure to exogenous mutagens^27^. Conversely, in subtypes with a lower mutational burden, it was the WGD^+^ tumors within that subtype with the higher ploidy-corrected mutational burden (Fig. 1b). This supports a recent report that predicts highly mutated tumors, which experience fewer somatic copy number alterations (SCNAs), encounter selection pressures that disfavor WGD, while tumor types with a lower mutational burden and increased SCNAs will favor WGD due to its capacity to buffer against deleterious mutations in genomic regions of loss of heterozygosity^10^.

We next explored the mutational landscape of WGD^+^ tumors, where we observed a significant enrichment of mutations in *TP53* and *PPPR21A* (Fig. 1c), consistent with findings from advanced cancer patients and a smaller cohort of TCGA samples^12,24^. The positive selection for these mutations is clear: p53 represents a major barrier to the proliferation of WGD^+^ cells, and thus inactivating mutations in *TP53* are favored in WGD^+^ cancers. Mutations in *PPP2R1A* promote centrosome clustering, an important adaptation for preventing multipolar cell division and cell death in WGD^+^ cells with supernumerary centrosomes^6,28^. We also identified mutations that are negatively enriched in WGD^+^ tumors, implying that these mutations are either less important for, or perhaps incompatible with, driving tumorigenesis in the context of WGD (Fig. 1c).

To assess changes in the microenvironment of WGD tumors, we applied the ABSOLUTE algorithm to infer the purity (*i.e.* the fraction of non-tumor cells) of TCGA tumor samples^23^. We found that WGD correlates with decreased purity and increased non-immune stromal infiltration (Extended Data Fig. 2a-b). We also assessed the correlation of WGD with TCGA estimates of tumor-infiltrating leukocytes (TILs) and found a negative correlation between WGD and TILs (Fig. 1d)^29,30^. When we performed gene expression analysis to identify genes differentially expressed in WGD^+^ tumors relative to WGD^−^ tumors, we found that the most negatively enriched gene sets in WGD^+^ tumors were inflammatory processes, further corroborating our finding that these tumors present with diminished host immune response similar to highly aneuploid tumors (Fig. 1e)^31,32^. We further identified that WGD^**+**^ tumors tend to overexpress genes important for cellular proliferation, mitotic spindle formation, and DNA repair (Fig. 1e, Supplementary Table 1). Collectively, our data demonstrate key genetic and phenotypic differences between WGD^+^ and WGD^−^ tumors, support the prognostic and therapeutic significance of WGD, and hint at potential adaptations and vulnerabilities that may inexorably arise following a WGD event.

## WGD confers unique genetic vulnerabilities on tumors

We examined whether WGD confers unique genetic dependencies on tumor cells by applying the ABSOLUTE algorithm to cancer cell lines from Project Achilles, which is a comprehensive catalog quantifying the essentiality of ∼20,000 genes across ∼600 cell lines following both CRISPR and RNAi-mediated gene depletion (Supplementary Table 2)^33-35^. After classifying the cell lines as either WGD^+^ or WGD^−^, we used Project Achilles data to score genes based upon their enrichment for essentiality in WGD^+^ cell lines relative to WGD^−^ cell lines (so-called ploidy-specific lethal (PSL) genes^21^) (Fig. 2a-b, Extended Data Fig. 2c, Supplementary Tables 3 and 4, *see methods for scoring details*). We mapped these PSL genes against the gene expression signature of WGD^**+**^ tumors and found several PSL genes to be significantly overexpressed, reinforcing their importance in the progression of WGD^**+**^ tumors (Fig. 2b-c).

**Figure 2.**
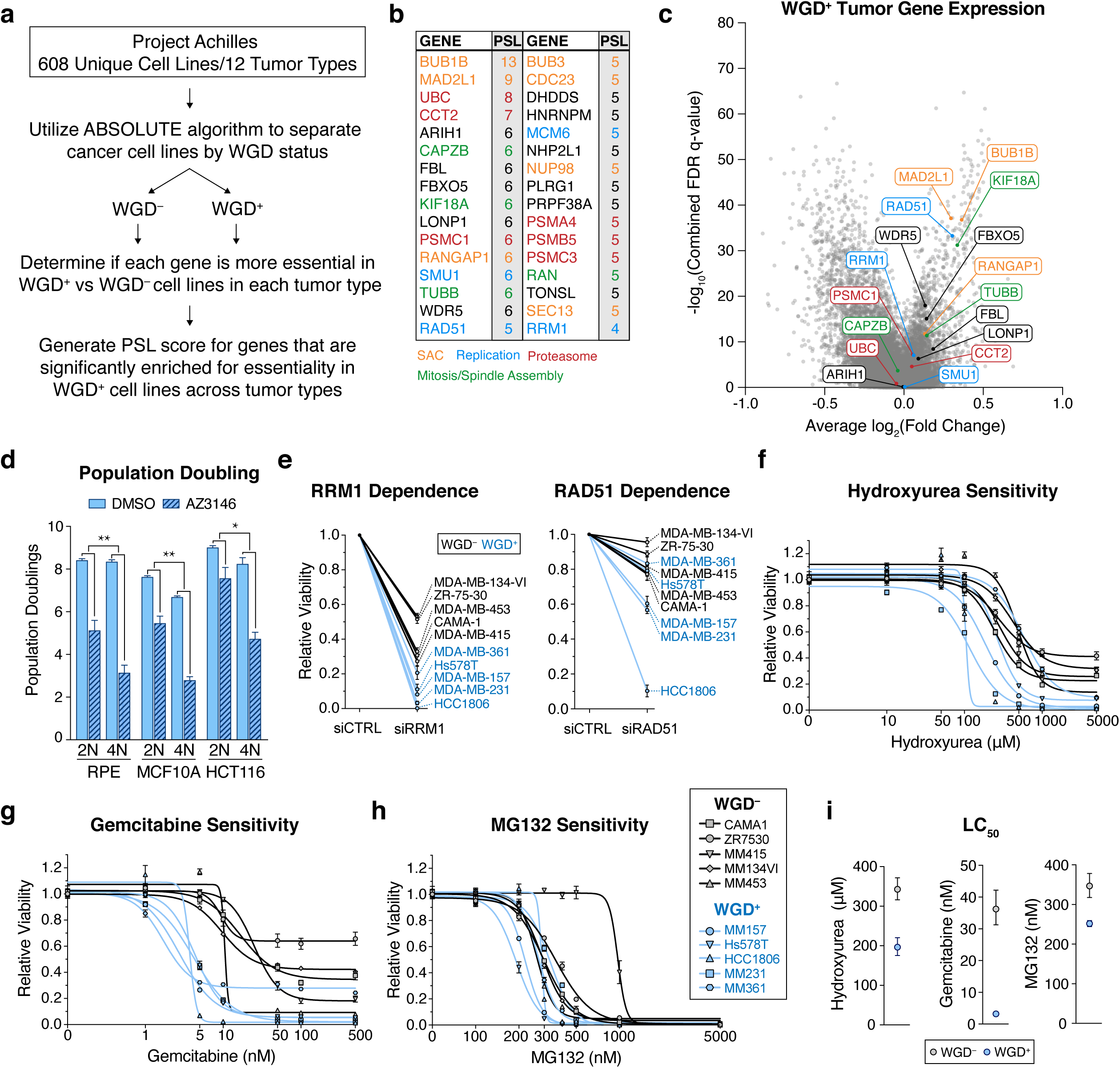
Identification and validation of PSL genes. **(a)** Workflow used to identify gene essentiality in WGD^+^ cancer cells from Project Achilles data *(see methods)*. **(b)** Top hits from PSL analysis; text color indicates genes associated with indicated pathways. **(c)** Gene expression fold changes in WGD^+^ tumors relative to WGD^−^ tumors plotted against combined FDR values across all tumor types with select PSL genes highlighted. **(d)** Population doublings after 8 days of AZ3146 treatment (two-way ANOVA with interaction; graph shows mean +/- SEM). **(e)** Relative viability of indicated cell lines 7 days after treatment with indicated siRNA (graph shows mean +/- SEM). **(f-h)** Dose-response curves for 5 WGD^−^ (black) and 5 WGD^+^ (blue) breast cancer cell lines 7 days after indicated drug treatment at the indicated concentrations (nonlinear regression with variable slope; graph shows mean +/- SEM at each dose). **(i)** Composite LC_50_ for 5 WGD^−^ and 5 WGD^+^ breast cancer cell lines for indicated drug treatments (nonlinear regression; graphs show LC_50_ +/- 95% CI). **siRNA Sequences** p < 0.05, ** p < 0.01, *** p < 0.001, **** p < 0.0001

To validate these PSL genes, we first generated three isogenically matched diploid (WGD^−^ or 2N) and tetraploid (WGD^+^ or 4N) cell lines as previously described (Extended Data Fig. 2d-g)^6,36^. These lines included the non-transformed epithelial cell lines RPE-1 and MCF10A, as well as the colon cancer cell line HCT116. Importantly, the development of these lines enabled us to directly compare cellular dependencies in cells differing only by WGD status.

We first validated *BUB1B* and *MAD2L1*, the two strongest PSL gene hits from our analysis. These genes encode proteins that are essential to the function of the spindle assembly checkpoint (SAC), which delays anaphase onset until all chromosomes have attached to the mitotic spindle, thus promoting the faithful partitioning of genomic content into two daughter cells during mitosis^37^. It has been demonstrated that increasing chromosome number prolongs the time needed to achieve full chromosome attachment and alignment^38^, suggesting that premature anaphase induced by disruption of the SAC should give rise to chromosome segregation errors at elevated rates in tetraploid cells^38^. Using live-cell imaging, we found that tetraploid cells indeed require more time to attach and align chromosomes relative to diploids in all three cell lines tested (Extended Data Fig. 3a). Consequently, we found that inhibition of the SAC using the small molecule inhibitor AZ3146, which inhibits the MPS1 kinase and abrogates the SAC in a manner similar to MAD2 or BUBR1 depletion, leads to a significant increase in chromosome segregation defects and micronuclei formation in tetraploid cells relative to diploids (Extended Data Fig. 3b). Micronuclei and chromosome segregation errors impair cell fitness, and concordantly, population doubling assays confirmed that tetraploid cells are significantly more sensitive to SAC inhibition than diploids (Fig. 2d). These data corroborate previous studies and served to validate our PSL analysis methodology^39,40^.

The identification of several genes involved in DNA replication as PSL hits suggests that WGD^+^ cells may also be more vulnerable to challenges to DNA replication than WGD^−^ cells. We first validated that reductions in the levels of RRM1 and RAD51 (two PSL genes known to mitigate the DNA damage associated with replication stress) preferentially impair the viability of tetraploid cells (Extended Data Fig. 3c-f). As an orthogonal approach, we also treated isogenic diploid and tetraploid cells with hydroxyurea or gemcitabine, which inhibit ribonucleotide reductase (RRM1) activity and induce replication stress. We observed that tetraploid cell lines show an increased sensitivity to these inhibitors relative to diploids (Extended Data Fig. 4a-b). We also confirmed this result in a panel of ten breast cancer cell lines (five WGD^+^ and five WGD^−^) (Fig. 2e-g,i, Extended Data Fig. 4d-e). These data reveal that WGD^+^ tumor cells are more dependent on specific DNA replication factors relative to WGD^−^ tumor cells, perhaps as a means to compensate for increased replication stress induced by tetraploidy^41,42^. These results are particularly significant in therapeutic contexts as gemcitabine and other inhibitors of ribonucleotide reductase represent the standard of care for treatment regimens across multiple cancer subtypes, and biomarkers that can predict sensitivity to gemcitabine hold real prognostic value^43^.

We also identified several PSL genes that encode for regulators of the proteasome, suggesting that WGD confers vulnerability to disruptions in protein stability/turnover. Indeed, we found that WGD^+^ cells are more sensitive to the proteasome inhibitor MG132 than WGD^−^ cells (Fig. 2h-i, Extended Data Fig. 4c). This dependency can likely be attributed to the highly aneuploid nature of WGD^+^ cells, as aneuploidy has previously been shown to induce proteotoxic stress ^44^. Supporting this view, we found that tetraploid RPE-1 cells, which maintain an euploid number of chromosomes (92) (Extended Data Fig. 2g), were the only cell line not more sensitive to MG132 relative to diploids (Extended Data Fig. 4c).

## WGD confers dependence on KIF18A

Our analysis identified the gene *KIF18A*, which encodes for a mitotic kinesin protein, as a significant PSL hit (Fig. 2b). KIF18A functions to suppress chromosomal oscillations at the metaphase plate by regulating microtubule dynamics to facilitate proper alignment and distribution of chromosomes during mitosis^45-48^. Importantly, in contrast to the aforementioned genes that regulate essential cellular processes such as SAC function, DNA replication, and proteasome activity, *KIF18A* is a non-essential gene in normal diploid cells, as attested by the fact that transgenic *KIF18A* knockout mice survive to adulthood^49,50^. Further, *KIF18A* is commonly overexpressed in WGD^+^ tumors (Fig. 2c). Its high PSL score and preferential gene expression in WGD^+^ tumors, combined with its dispensability in normal diploid cells, make *KIF18A* an exciting new candidate for therapeutic exploration.

We first validated *KIF18A* as a PSL gene by confirming that depletion of KIF18A significantly impairs the viability of tetraploid but not diploid cells (Fig. 3a, Extended Data Fig. 5a). To understand the mechanism underlying this reduction in viability, we used live-cell imaging to monitor mitotic progression following KIF18A depletion in our three isogenic diploid and tetraploid cell models. This analysis revealed that KIF18A knockdown has profoundly differential effects on the fidelity of mitosis in tetraploid cells relative to diploid cells. We observed that depletion of KIF18A had no effect on mitotic duration in diploid cells. By contrast, depletion of KIF18A led to significantly prolonged mitoses in tetraploid cells (Fig. 3b). We also observed that while diploid cells lacking KIF18A exhibited subtle defects in chromosome misalignment at anaphase onset, chromosome segregation proceeded relatively normally with no significant increase in the generation of micronuclei following mitosis (Fig. 3b,g). By contrast, tetraploid cells depleted of KIF18A exhibited significant increases in chromosome misalignment, anaphase lagging chromosomes, and micronuclei formation (Fig. 3b,g Extended Data Fig. 5b,f, Extended Data Fig. 6a, Supplementary Movies 1-4).

**Figure 3.**
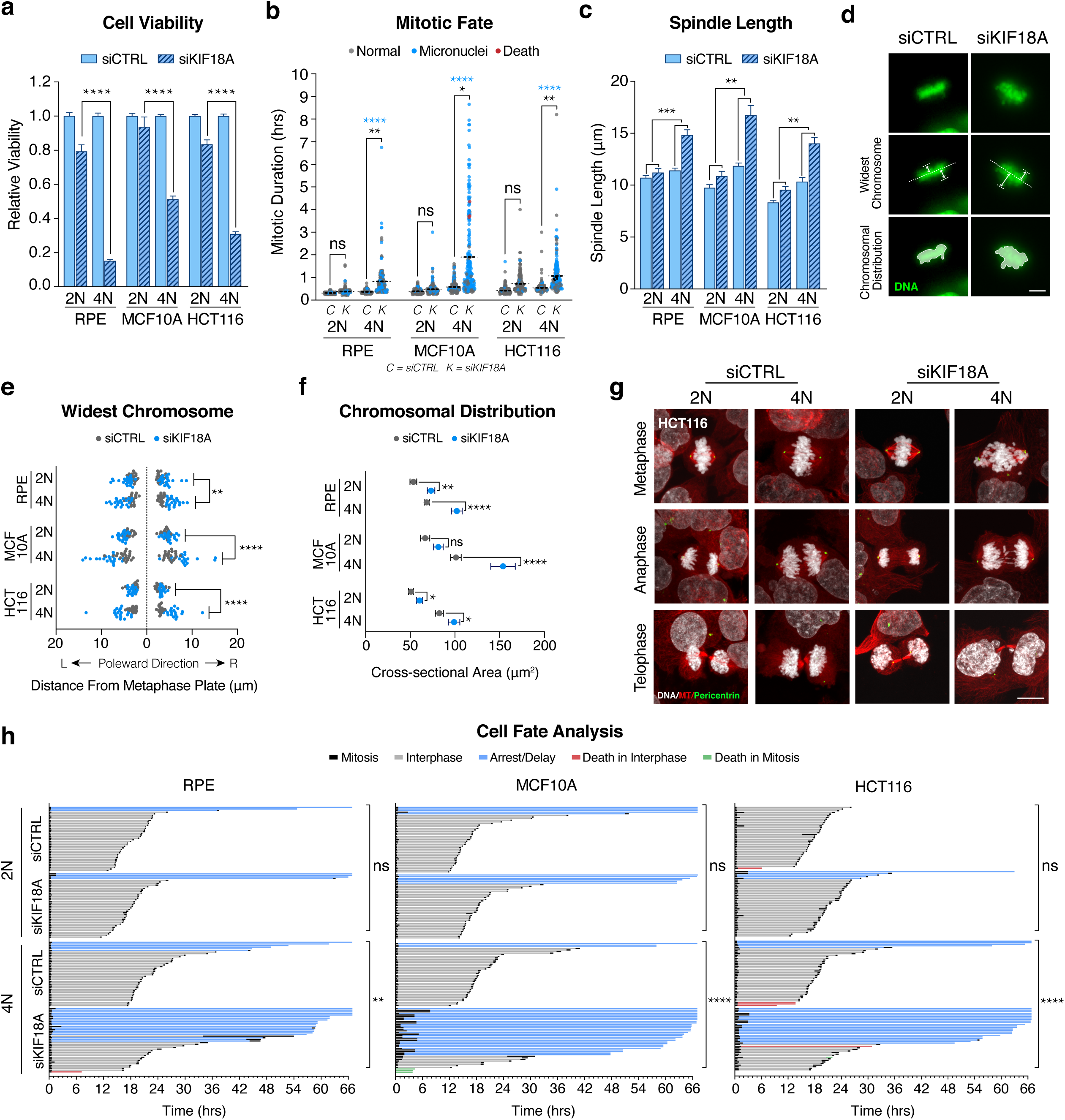
KIF18A depletion impairs the mitotic fidelity of WGD^+^ cells. **(a)** Relative viability of indicated cell lines 8 days after transfection with the indicated siRNAs (each condition normalized to respective control; Student’s unpaired t-test; graph shows mean +/- SEM). **(b)** Mitotic duration and fate after treatment with indicated siRNA (n = 200 cells per condition; black stars indicate p-value for Student’s t-test comparing mean mitotic duration; blue stars indicate p-value for Fisher’s exact test comparing the fraction of mitoses that give rise to micronuclei; dotted line represents mean mitotic duration). **(c)** Measurement of spindle length (centrosome-to-centrosome) after transfection with indicated siRNA (n = 20 cells per condition; two-way ANOVA with interaction; graph shows mean +/- SEM; scale bar 10 μm). **(d)** Image demonstrating how we measured chromosome oscillations immediately prior to anaphase by assessing the widest oscillating chromosomes in each poleward direction and the cross-sectional area of all the chromosomes. **(e)** Widest oscillating chromosome in each poleward direction immediately prior to anaphase (n = 20 cells per condition; two-way ANOVA with interaction). **(f)** Two-dimensional cross-sectional area of the entire body of chromosomes immediately prior to anaphase (n = 20 cells per condition; Student’s unpaired t-test; graph shows mean +/- SEM). **(g)** Representative confocal images showing phases of mitosis in indicated cell lines 48 hours after transfection with indicated siRNA (scale bar 10 μm). **(h)** Cell fates of indicated cell lines tracked for 3 days beginning 18 hours after transfection with indicated siRNA (n = 40 cells per condition; Fisher’s exact test comparing fraction of cells arresting/delaying in interphase relative to control group). p < 0.05, p < 0.01, p < 0.001, p < 0.0001

It has been demonstrated that the nuclear membranes surrounding micronuclei are prone to rupture, thereby exposing the chromosomal contents harbored within the micronuclei to the cytosolic environment^51^. This defect induces both catastrophic DNA damage to the exposed chromosomes as well as stimulation of the cGAS-STING pathway^52-54^. Indeed, we found that micronuclei in cells depleted of KIF18A showed both γ-H2AX and cGAS labeling (Extended Data Fig. 5d). Of note, we observed that a greater fraction of micronuclei in tetraploid cells are cGAS^+^ compared to diploid cells, and a greater fraction of micronuclei arising in tetraploid cells depleted of KIF18A are cGAS^+^ compared to micronuclei induced by SAC impairment (Extended Data Fig. 5d). These data indicate that tetraploid cells depleted of KIF18A give rise to micronuclei that are particularly fragile and prone to rupture, a characteristic that likely contributes to the observed differential effect on viability.

We speculated that the mitotic delays and aberrant chromosome segregation defects observed following KIF18A loss may be induced by changes in spindle morphology in tetraploid cells. To accommodate their doubled chromosome content, tetraploid cells assemble larger mitotic spindles^6^. Indeed, we found that spindles in tetraploid cells were on average ∼17% longer than in diploids (Fig. 3c). Depletion of KIF18A led to an additional increase in spindle length, and this effect was significantly more dramatic in tetraploid cells relative to diploids (Fig. 3c).

We also measured the magnitude of chromosome oscillations immediately prior to anaphase onset in diploid and tetraploid cells by assessing the widest oscillating chromosomes in each poleward direction, as well as the overall chromosome alignment efficiency by measuring the total two-dimensional area occupied by the entire body of chromosomes (Fig. 3d). These analyses revealed that the magnitude of chromosomal oscillations is significantly greater in tetraploid cells relative to diploid cells following KIF18A depletion (Fig. 3e-f). One consequence of hyper-oscillating chromosomes in tetraploid cells depleted of KIF18A is that they have a propensity to lose their attachment to the mitotic spindle and activate the spindle assembly checkpoint, thus explaining the mitotic delays we observed (Extended Data Fig. 6b)^55,56^. A second consequence is that severely misaligned chromosomes must traverse a significantly greater distance during anaphase in tetraploid cells compared to diploid cells, thus explaining the observed increase in lagging chromosomes and micronuclei.

Numerous studies have indicated that aneuploidy and micronuclei induced by lagging chromosomes can impair cell proliferation, in part through activation of the p53 pathway^17^. We therefore used long-term live-cell imaging to track the fates of isogenic diploid and tetraploid cells depleted of KIF18A. Our analysis revealed that while the majority of diploid cells depleted of KIF18A undergo normal cell cycle progression, isogenic tetraploid cells depleted of KIF18A are prone to interphase cell cycle arrest following abnormal mitosis, concomitant with p53 pathway activation (Fig 3h and Extended Data Fig. 5e). Thus, our data reveal that loss of KIF18A in WGD^+^ cells predisposes cells to lagging chromosomes, micronuclei formation, micronuclei rupture, and proliferative arrest. Supporting this mechanism, we found that cellular proliferation is required for the loss of KIF18A to drive our observed viability defects (Extended Data Fig. 5c).

We sought to also validate the ploidy-specific lethality of KIF18A across our panel of breast cancer cell lines. Supporting our pan-cancer gene expression analysis (Fig. 2c), we found that KIF18A protein levels are typically elevated in WGD^+^ cells (Fig. 4a, Extended Data Fig. 7a). Knockdown of KIF18A from all ten breast cancer cell lines (Extended Data Fig. 7b) confirmed that WGD^+^ breast cell lines experience a significantly greater reduction in viability relative to WGD^−^ cell lines (Fig. 4b, Extended Data Fig. 7c-e, Supplementary Movies 5-6). Live-cell imaging revealed that WGD^+^ breast cancer cells exhibited increased spindle lengths and chromosome hyper-oscillations relative to WGD^−^ breast cancer cells after loss of KIF18A (Fig. 4d-e, Extended Data Fig. 7f), thus promoting chromosome detachment, spindle assembly checkpoint activation, and prolonged mitosis (Fig. 4c, Extended Fig. 6c). Notably, we observed that a large fraction of WGD^+^ cells were never able to satisfy the spindle assembly checkpoint and exhibited a dramatically prolonged mitotic arrest before ultimately undergoing mitotic cell death (Fig. 4c). WGD^+^ cells depleted of KIF18A that were able to achieve anaphase exhibited significant increases in both anaphase lagging chromosomes and micronuclei relative to the WGD^−^ cell lines, similar to what was observed in the isogenic tetraploid models (Fig. 4c, Extended Fig. 7g). However, in contrast to the p53-proficient isogenic tetraploid cells, WGD^+^ breast cancer cell lines depleted of KIF18A were not prone to cell cycle arrest following abnormal mitosis, likely due to the fact that all WGD^+^ lines have impaired p53 function (Fig. 4f). Instead, a fraction of these cells die in interphase after experiencing catastrophic mitoses resulting in micronuclei formation, while the majority of these WGD^+^ cells initiate a second round of mitosis without KIF18A, where they are just as likely or more prone to mitotic cell death (Fig. 4g).

**Figure 4.**
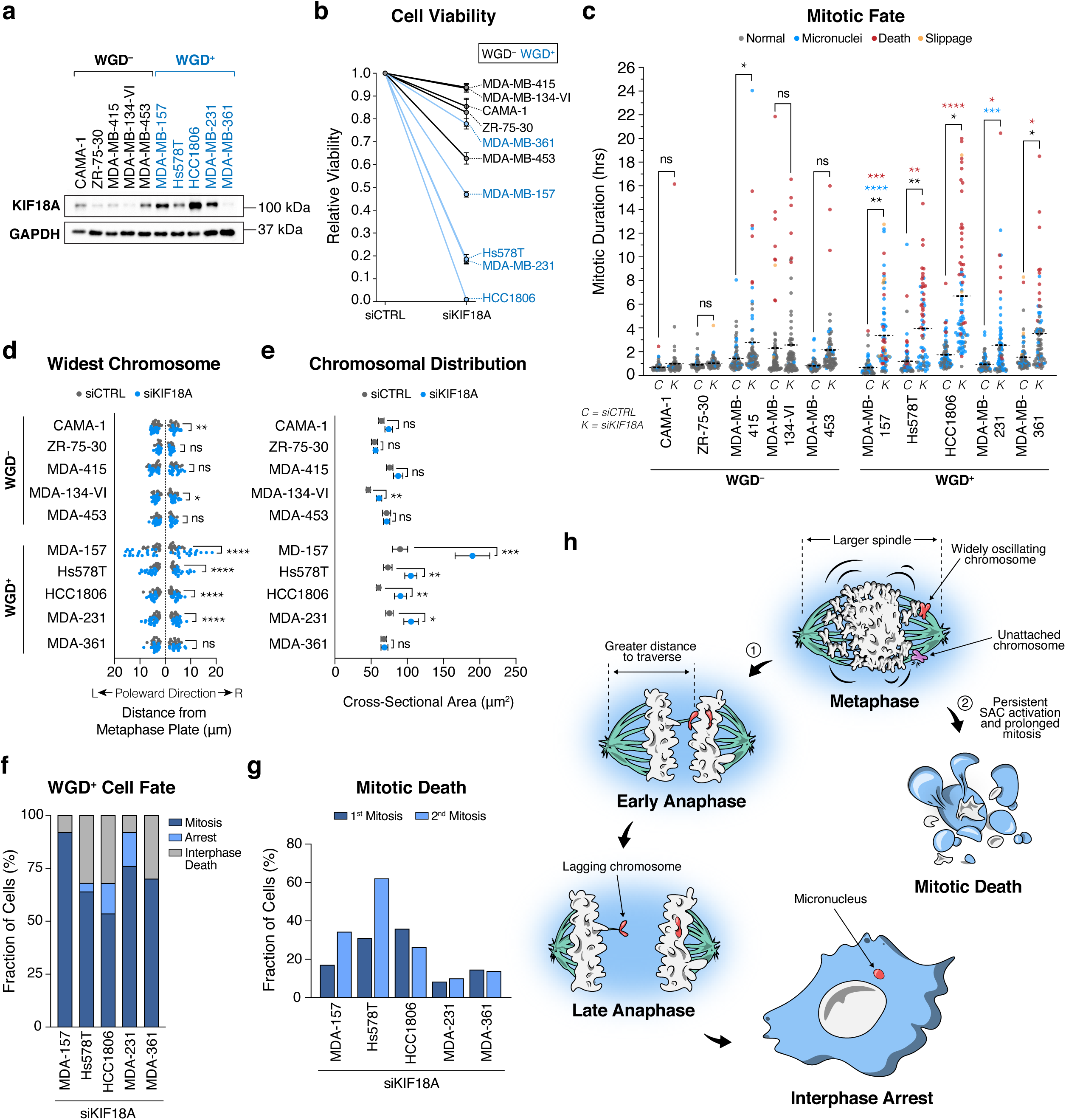
WGD confers dependence on KIF18A in a panel of breast cancer cell lines. **(a)** Western blot showing endogenous KIF18A levels in indicated cell lines. **(b)** Relative viability of cell lines 8 days after transfection with the indicated siRNAs. **(c)** Mitotic duration and fate following transfection with indicated siRNA (dotted line represents mean mitotic duration; black stars indicate p-values for Student’s unpaired t-test comparing mean mitotic duration; blue stars indicate p-values for Fisher’s exact test comparing fraction of mitoses that give rise to micronuclei; red stars indicate p-values for Fisher’s exact test comparing fraction of cell that die in mitosis; n = 80 cells per condition). **(d)** Widest oscillating chromosome in each poleward direction immediately prior to anaphase (n = 20 cells per condition; Student’s unpaired t-test). **(e)** Two-dimensional cross-sectional area of the entire body of chromosomes immediately prior to anaphase (n = 20 cells per condition; Student’s unpaired t-test; graph shows mean +/- SEM). **(f)** The fraction of cells in each cell line that undergo indicated fates after completing a mitosis deficient of KIF18A that resulted in micronuclei formation (n = 25 cells per condition). **(g)** The fraction of cells in each cell line that experience mitotic death in their first and second mitoses following KIF18A depletion (n = 25 cells per condition). **(h)** Depletion of KIF18A impairs WGD^+^ cell viability through two mechanisms: 1 - larger spindles and wider oscillations increase the distance some chromosomes must traverse in anaphase leading to lagging chromosomes, micronuclei formation, and cellular arrest. 2 - widely oscillating chromosomes fail to properly attach to microtubules, thus activating the spindle assembly checkpoint and leading to prolonged mitosis and death. * p < 0.05, ** p < 0.01, *** p < 0.001, **** p < 0.0001

Collectively, these data reveal that loss of KIF18A specifically impairs mitotic fidelity and cell viability in WGD^+^ cancer cells (Fig. 4f), highlighting KIF18A as an attractive new therapeutic target whose inhibition may enable the specific targeting of WGD^**+**^ tumors while sparing the normal diploid cells that comprise human tissue. Supporting this view, it has been demonstrated that *KIF18A* knockout mice are protected from colitis-associated colorectal tumors and that depletion of KIF18A from the WGD^+^ breast cancer cell line MDA-MB-231 impairs tumor growth *in vivo*^57,58^.

An important consideration is that WGD^+^ cancer cells exhibit numerous characteristics that distinguish them from WGD^−^ cells in addition to simply having extra chromosomes and larger spindles: WGD^+^ cells frequently possess supernumerary centrosomes, are chromosomally unstable, and have inactivating mutations in *TP53*^6,14^. However, we favor a model in which the dependency of WGD^+^ cells on KIF18A is due predominantly to the extra chromosomes, as we observe viability defects in tetraploid RPE-1 cells despite the fact that they possess a euploid complement of chromosomes, are chromosomally stable, and have functional p53 signaling.

Nevertheless, we do note that some WGD^−^ cancer lines show sensitivity to KIF18A depletion, suggesting that other defects may exist that predispose to KIF18A sensitivity. Indeed, Marquis et al., (unpublished) propose that altered spindle microtubule dynamics in chromosomally unstable cancer cells may also induce KIF18A sensitivity.

Herein, we have comprehensively catalogued specific genomic characteristics unique to WGD^**+**^ tumors and demonstrated that WGD confers specific, exploitable vulnerabilities on tumor cells. It should be noted that highly aneuploid cancer cells (*e.g.* possessing ≥ triploid number of chromosomes) almost exclusively arise from WGD^+^ cells that have lost chromosomes over many cell divisions (Fig. 1a)^21,22^. By contrast, WGD^−^ tumors, which are also typically aneuploid but maintain a chromosome number in the near-diploid range, do not exhibit the same level of dependencies. This suggests that aneuploidy *per se* is insufficient to drive the dependencies we observe, but rather it is the overall increase in chromosome number that is critical.

Our combined computational and *in vitro* approaches have further characterized the genetic landscape of WGD^+^ tumors and generated a list of ploidy-specific lethal (PSL) genes that highlight the vulnerabilities that can arise with a WGD event. We have also identified a new therapeutic target in KIF18A, which holds the potential of broad applicability with minimal toxicity. Collectively, this work serves to underscore the importance and untapped potential of exploring and targeting WGD in human tumors.

## Acknowledgements

The results published here are in part based upon data generated by the TCGA Research Network: https://www.cancer.gov/tcga. We would like to thank Janice Weinberg for statistical advice, Jason Stumpff for technical advice, reagents, and sharing unpublished data, David Pellman for sharing reagents, and Scott Carter for sharing ABSOLUTE data for cell lines in the CCLE. R.J.Q. is supported by a Canadian Institutes of Health Research Doctoral Foreign Study Award. N.J.G. is a member of the Shamim and Ashraf Dahod Breast Cancer Research Laboratories and is supported by NIH grants CA154531 and GM117150, the Karin Grunebaum Foundation, the Smith Family Awards Program, the Melanoma Research Alliance, and the Searle Scholars Program. This work was also supported in part by a pilot grant from the ACS and the Boston University Clinical and Translational Science Institute Bioinformatics Group, who are supported by a grant from the NIH/NCATS (1UL1TR001430).

## Author Contributions

RQ and NG designed the experiments and wrote the manuscript. RQ performed the cell biological assays and imaging analysis. AD, CT, and SP assisted RQ with the cell biological assays. KK and MV performed imaging analysis. TK generated the isogenic diploid and tetraploid HCT-116 cells. JV generated cell lines. NH and AM performed animal studies. AT, YK, NP, MM, and JC performed the computational analyses. All authors edited the manuscript.

## Supplementary Tables

**Supplementary Table 1**

Pan-cancer gene expression analysis with GSEA and gene expression analysis for each tumor subtype for WGD^+^ tumors in the TCGA.

**Supplementary Table 2**

ABSOLUTE algorithm applied to cancer cell lines indicating purity, ploidy, and number of whole genome doublings.

**Supplementary Table 3**

Gene essentiality data for ∼600 cancer cell lines from the Cancer Cell Line Encyclopedia showing genes enriched for essentiality in WGD^+^ cell lines.

**Supplementary Table 4**

List of ploidy-specific lethal genes ranked by their PSL score.

## Supplementary Movies

**Supplementary Movie 1**

Live-cell imaging of diploid (2N) MCF10A H2B-GFP cells following transfection with control siRNA (5 frames/second; hour:minute; scale bar 10μm).

**Supplementary Movie 2**

Live-cell imaging of diploid (2N) MCF10A H2B-GFP cells following transfection with KIF18A siRNA (5 frames/second; hour:minute; scale bar 10μm).

**Supplementary Movie 3**

Live-cell imaging of tetraploid (4N) MCF10A H2B-GFP cells following transfection with control siRNA (5 frames/second; hour:minute; scale bar 10μm).

**Supplementary Movie 4**

Live-cell imaging of tetraploid (4N) MCF10A H2B-GFP cells following transfection with KIF18A siRNA (5 frames/second; hour:minute; scale bar 10μm).

**Supplementary Movie 5**

Live-cell imaging of the HCC1806 H2B-GFP breast cancer cell line following transfection with control siRNA (40 frames/second; hour:minute; scale bar 100μm).

**Supplementary Movie 6**

Live-cell imaging of the HCC1806 H2B-GFP breast cancer cell line following transfection with KIF18A siRNA (40 frames/second; hour:minute; scale bar 100μm).

## Methods

### WGD/Purity/Ploidy Calls

TCGA samples were previously analyzed using the ABSOLUTE algorithm^4^. ABSOLUTE takes copy number and mutation data to estimate sample purity, ploidy, and number of whole genome doublings. ABSOLUTE calls for TCGA samples are available in ref^5^. Briefly, the algorithm infers from sequencing data what fraction of a tumor sample is composed of tumor cells vs non-tumor cells (purity) as well as the ploidy of a tumor sample by analyzing copy number ratios across the entire genome. WGD status is inferred based on the ploidy distribution within a tumor type, the homologous copy number information across the genome, and the presence of duplicated mutations.

### Ploidy-Corrected Mutational Burden

To compare the ploidy-corrected mutational burden of WGD^+^ and WGD^−^ TCGA samples, we divided the non-synonymous mutations per Mb (log10 transformed)^1^ of each sample by their ploidy as defined by ABSOLUTE. We performed a linear regression using the lm function in R version 3.2.3. The formula was:

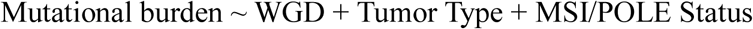

We applied the Wilcoxon rank-sum test to analyze the total mutational burden and ploidy-corrected mutational burden between WGD^+^ and WGD^−^ samples within each subtype.

### Mutations in WGD^+^ Tumors

To identify gene mutational frequencies associated with WGD status, we applied logistic regression to 631 driver genes that were found to be significantly recurrently mutated in one or more tumor types by MutSig2CV^2^. The formula for the logistic regression model was:

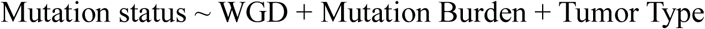

where Mutation Burden was the number of non-synonymous mutations per Mb (log10 transformed)^1^ and WGD status was defined by ABSOLUTE calls retrieved from http://api.gdc.cancer.gov/data/4f277128-f793-4354-a13d-30cc7fe9f6b5. The maf file from TCGA PanCanAtlas MC3^3^ project was used to derive the mutation status for each gene in each tumor retrieved from https://api.gdc.cancer.gov/data/1c8cfe5f-e52d-41ba-94da-f15ea1337efc. This file was filtered to only include variants with “PASS”, “wga”, or “native_wga_mix” in the “FILTER” column. Variants with “Frame_Shift_Del”, “Frame_Shift_Ins”, “In_Frame_Del”, “In_Frame_Ins”, “Missense_Mutation”, “Nonsense_Mutation”, “Nonstop_Mutation”, “Translation_Start_Site”, “Splice_Site”, “De_novo_Start_InFrame”, “De_novo_Start_OutOfFrame”, “Stop_Codon_Del”, “Stop_Codon_Ins”, “Start_Codon_Del”, or “Start_Codon_Ins” in the “Variant_Classification” column were considered non-synonymous. An FDR correction was applied to the p-values for the WGD term to control for multiple hypothesis testing.

### Leukocyte infiltrate and stromal calls

Estimates of leukocyte fraction in the TCGA samples were generated using a mixture model of DNA methylation in pure leukocytes versus normal tissue. More details and all calls can be found in ref^6^. Stromal calls were made by subtracting leukocyte fraction from ABSOLUTE purity estimates described above. Spearman correlation coefficients were calculated after removing MSI/POLE mutant samples from the dataset and using the spearmanr function using cor.test in R (method = “spearman”), which was run using R version 3.2.3.

### Gene Expression Analysis

Expression and copy number data of TCGA samples were obtained from the PanCanAtlas project (https://gdc.cancer.gov/about-data/publications/pancanatlas). RNA-seqV2 data was used for expression analysis (http://api.gdc.cancer.gov/data/3586c0da-64d0-4b74-a449-5ff4d9136611). Expression values were log2-transformed after adding a pseudo-count of 1. Copy number ratios were obtained for each gene by running GISTIC2.0 on the PanCan segmentation file (http://api.gdc.cancer.gov/data/00a32f7a-c85f-4f86-850d-be53973cbc4d). Analysis was limited to primary tumors across all cancer types. P-values for WGD were corrected for multiple hypothesis testing with the Benjamini-Hochberg False Discovery Rate (FDR).

To identify gene expression profiles associated with WGD status, we applied the following linear model to each gene within each tumor type:

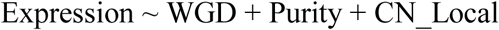

where Purity is the ABSOLUTE-estimated purity for each tumor and CN_Local is the log2 copy number ratio for that gene in each tumor estimated by GISTIC2.0 [ref: 21527027].

Note that the CN_Local variable was different for each gene (as each gene has a different copy number profile) while the WGD and Purity variables were the same for all genes. The Benjamini Hochberg method was used to correct p-values from the WGD term for multiple hypothesis testing. Genes were considered significantly associated with WGD status if they had an FDR q-value less than 0.05. Genes up-regulated in more than 10 tumor types were analyzed with hypeR [ref: 31498385] using the MSigDB Hallmark gene sets to identify biological categories enriched among these genes. Similarly, genes down-regulated in more than 10 tumor types were also analyzed with hypeR in the same fashion. To generate a volcano plot across tumor types, the coefficient for WGD was averaged and the FDR-corrected q-values were combined using the Fisher’s method.

### Ploidy-specific lethal (PSL) score analysis

#### Thresholded Analysis

Genes were assigned a binary classification (essential or non-essential) based on cutoffs established by Project Achilles. In the database, a score of -1 is assigned to a gene when its depletion in a given cell line results in a viability defect equal to the depletion of a curated list of gold standard common-essential genes^7,8^. Based on this scoring system, we defined any gene with a score ≤ -1 for a given cell line as essential. We then compared the fraction of cell lines in the WGD^−^ and WGD^+^ groups where a gene was essential. When a gene was essential in a significantly greater fraction of WGD^+^ cell lines than WGD^−^ cell lines (Fisher’s exact test, p < 0.1) in a specific tumor subtype, it was considered a “hit” in this analysis (Extended Data Fig. 3a).

#### Non-thresholded analysis

Within each tumor type, the median essentiality scores for each gene in the WGD^−^ and WGD^+^ cell lines were identified. When a gene showed a statistically significant enrichment in its median essentiality score in the WGD^+^ compared to the WGD^−^ cell lines (Wilcoxon test, p < 0.05), and also had an essentiality score of ≤ -0.5 in the WGD^+^ cell lines, it was considered at “hit” in this analysis (Extended Data Fig. 3a).

#### Final PSL score

We employed the thresholded analysis with the Fisher’s exact test and non-thresholded analysis with the Wilcoxon rank-sum in each individual tumor type (n=12) as well as in a combined pan-cancer analysis. These analyses were also performed separately for the CRISPR and RNAi datasets. Only genes that had measurable data in 95% of total cell lines were analyzed. The final PSL score for each gene was the total number of instances a gene was found to be a hit across all analyses (Fig. 2b, Supplementary Table 3,4). As a result, some hits may have come entirely from either the CRISPR or RNAi datasets, such as KIF18A which was only found to be enriched for essentiality in the CRISPR dataset, likely due to insufficient knockdown in the RNAi dataset.

### Cell Culture

All breast cancer cell lines were purchased from ATCC and used at early passage numbers. Isogenic tetraploid cell lines were generated as described^6^. hTERT-RPE-1 were cultured in DME/F12 (HyClone) supplemented with 10% FBS (ThermoFisher) with 50□IU/mL penicillin and 50 µg/mL streptomycin (ThermoFisher). HCT116, CAMA-1, MDA-MB-415, MDA-MB-134-VI, MDA-MB-157, Hs578T, MDA-MB-231, MDA-MB-361 cells were cultured in high glucose DMEM (Gibco) supplemented with 10% FBS with 50□IU/mL penicillin and 50□µg/mL streptomycin. ZR-75-30 and HCC1806 cells were cultured in RPMI (Gibco) supplemented with 10% FBS with 50□IU/mL penicillin and 50 µg/mL streptomycin. MCF10A cells were cultured in DME/F12 (HyClone) supplemented with 5% horse serum (ThermoFisher), 20ng/mL EGF (ThermoFisher), 500ng/mL hydrocortisone (ThermoFisher), 100ng/mL cholera toxin (Sigma), 10ug/ml insulin (ThermoFisher), with 50□IU/mL penicillin and 50 µg/mL streptomycin.

### siRNA Transfections

siRNA transfections using Lipofectamine RNAiMAX (Invitrogen) were performed according to the manufacturer’s instructions. The final concentration of KIF18A of CTRL siRNA in the medium was 10 nM, excepting MCF10A KIF18A siRNA transfections, which were performed at a final concentration of 1nM, and RRM1/RAD51 siRNA transfections, which were performed at a final concentration of 50 pM with CTRL siRNA adjusted accordingly.

### siRNA Sequences

Non-targeting control (CTRL) (Dharmacon) 5’-UGGUUUACAUGUCGACUAA-3’ KIF18A (Silencer Select s37882 – Ambion)^8^ 5’-UCUCGAUUCUGGAACAAGCAG-3’ RAD51 (Silencer Select s11735 – Ambion) 5’-UGAUUAGUGAUUACCACUGCT-3’ RRM1 (On-Target plus SMARTpool – Dharmacon)

5’-UAUGAGGGCUCUCCAGUUA-3’

5’-UGAGAGAGGUGCUUUCAUU-3’

5’-UGGAAGACCUCUAUAACUA-3’

5’-CUACUAAGCACCCUGACUA-3’

### Inducible shRNA

We infected cells with a SMARTvector Inducible Lentiviral shRNA (Horizon) targeting KIF18A and selected cells with puromycin (Santa Cruz Biotechnology) at 2 μg/mL. Cells were induced with doxycycline (Sigma) at 1 μg/mL for 7 days and viability was assessed.

shRNA sequence: 5’ **-**CGATGACACACATATAACACT-3’

### Inducible CRISPR-Cas9

We infected cells with pCW-Cas9 plasmid (Addgene #50661) and selected cells with puromycin at 2 μg/mL. To improve knockout efficiency, cells were then infected with 2 distinct KIF18A sgRNA plasmids. Each sgRNA sequence was cloned into its own lenti-sgRNA-blast plasmid (Addgene #104993) and these plasmids were co-packaged into lentivirus and used to infect cells, which were then selected with blasticidin (Sigma) at 5μg/mL. The sequences for both KIF18A targeting sgRNAs are available in ref^10^:

### Cell Viability Experiments

All cell viability assays were done using CellTiter-Glo (Promega) and performed according to the manufacturer’s instructions.

### Drug Treatments

AZ3146 (Tocris) was used at a concentration of 1µM in HCT116 cells, 2µM in MCF10A cells, and 4µM in RPE-1 cells. These concentrations were experimentally determined to be the minimum concentration required to inhibit the SAC in each respective cell line.

MG132 (Selleck Chemicals) was used at indicated concentrations.

### Antibodies

Rabbit polyclonal anti-KIF18A (Bethyl Cat # A301-080A)

Rabbit monoclonal anti-RRM1 (Cell Signaling Technology Cat # 8637)

Rabbit polyclonal anti-RAD51 (Santa Cruz Biotechnology Cat # sc-8349)

Rabbit monoclonal anti-cGAS (Cell Signaling Technology Cat # 15102)

Mouse monoclonal anti-phospho-histone H2A.X (Ser 139) (Sigma-Aldrich Cat # 05-636-I)

Mouse monoclonal anti-p53 (Santa Cruz Biotechnology Cat # sc-126)

Rabbit monoclonal anti-p21 (Cell Signaling Technology Cat # 2947)

Rabbit monoclonal anti-Cas9 (Active Motif Cat # 61978)

Rabbit monoclonal anti-GAPDH (Cell Signaling Technology Cat # 2118)

Mouse monoclonal anti-Vinculin (Abcam Cat # ab18058)

Mouse monoclonal anti-Tubulin (clone DM1A) (Sigma-Aldrich Cat # 05-829)

Rabbit polyclonal anti-Pericentrin (Abcam Cat # ab4448)

### Population Doubling Assay

10,000 cells were seeded in a 10cm dish with AZ3146 at indicated concentrations. Fresh drug was added every 3 days. After 8 days cells were counted, and population doublings were calculated using the formula 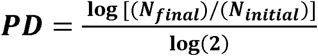

### Live-Cell Imaging

Stably expressing H2B-GFP cells were grown on glass-bottom 12-well tissue culture dishes (Cellvis) and treated with drugs or transfected with siRNAs of interest. At 24 hours post-treatment, imaging was performed on a Nikon TE2000-E2 inverted microscope equipped with the Nikon Perfect Focus system. The microscope was enclosed within a temperature and atmosphere-controlled environment at 37 °C and 5% humidified CO_2_. Fluorescent images were captured every 3 minutes with a 20X 0.5 NA Plan Fluor objective at multiple points for 72 hours. Captured images were analyzed for mitotic defects using NIS elements software.

### Chromosome Alignment Measurement

Live cell imaging was used to track H2B-GFP expressing cells to the frame immediately preceding anaphase and the distance from the metaphase plate to the widest oscillating chromosomes in each poleward direction was measured manually. We also measured the total chromosomal distribution immediately prior to anaphase by recording the area of automatically generated regions of interest (ROIs) based on fluorescence intensity using NIS elements software.

### Cell Fate Analysis

Live cell imaging was used to track cells treated with control siRNA to obtain the average cell cycle time for each cell line, and cells treated with siKIF18A were called as “arrested/delayed” if they spent greater than 3 standard deviations above the mean cell cycle time of control cells in interphase.

### Immunofluorescence Microscopy

Cells were plated on glass cover slips and then washed in microtubule stabilizing buffer (MTSB) (4M Glycerol, 100mM PIPES, pH 6.9, 1mM EGTA, 5mM MgCl_2_) for 1 min, extracted in MTSB-0.5% Triton for 2 min, and washed again in MTSB for 2 min. Cells were then fixed in 1% EM grade glutaraldehyde for 10 min. Glutaraldehyde was quenched by washing twice in NaBH_4_ in water for 12 min each. Cells were then blocked for 30 min in TBS-BSA (10 mM Tris, pH 7.5, 150 mM NaCl, 5% BSA, 0.2% sodium azide), and incubated with primary antibodies diluted in TBS-BSA for 60 min in a humidified chamber. Primary antibodies were visualized using species-specific fluorescent secondary antibodies (Molecular Probes) and DNA was detected with 2.5 µg/ml Hoechst. Confocal immunofluorescence images were collected at 405, 488, and 561 nm on a Nikon Ti-E inverted microscope with C2+ laser scanning head. A series of 0.5 µm optical sections were acquired using a 60x objective lens. Images presented in figures are maximum intensity projections of entire z-stacks.

### Spindle Length

Spindles were measured using immunofluorescence microscopy. Cells were stained for tubulin/centrosomes and spindle length was assessed by measuring the distance from centrosome to centrosome of cells in metaphase using NIS elements software.

### Western Blotting

Cells were rinsed with ice-cold 1X PBS (Boston Bioproducts) and lysed immediately with cell lysis buffer (2% w/v SDS, 10% Glycerol, 60 mM Tris-HCl) supplemented with 1X HALT protease and phosphatase dual inhibitor cocktail (ThermoFisher). Cell lysates were then sonicated for 15 seconds at 20 kHz and Sample Buffer (Boston Bioproducts) was added to a final concentration of 1X, after which protein samples were incubated at 95°C for 5 minutes.

Cell lysates were resolved via SDS-PAGE (Resolving/Separating gel: 7.5% acrylamide, 375 mM Tris-HCl (pH 8.8), 0.1% SDS, 0.25% ammonium persulfate, 0.15% tetramethylethylenediamine; Stacking gel: 4% acrylamide, 125 mM Tris-HCl (pH 6.8), 0.1% SDS, 0.5% ammonium persulfate, 0.3% tetramethylethylenediamine) in SDS-PAGE running buffer (25 mM Tris-HCl, 192 mM Glycine, 0.1% SDS). Samples were passed through the stacking gel layer at 130 V for 15 minutes, followed by resolution of samples at 230 V for 25 minutes. Samples were transferred to 0.45μm Immobilon PVDF membranes (EMD Millipore) using a wet-tank transfer system (Bio-Rad) in Towbin transfer buffer (25 mM Tris-HCl, 192 mM Glycine, 10% methanol) for 16 hours at 30 mA at 4°C. Following transfer, membranes were blocked in TBS-0.5% Tween-20 (10 mM Tris-HCl, 150 mM NaCl, 0.5% Tween-20) containing 5% non-fat dried milk (NFDM) for 1 hour, and then incubated with primary antibodies diluted in 1% NFDM TBS-0.5% Tween-20 solution. Membranes were rinsed in TBS-0.5% Tween-20 solution following primary and secondary antibody incubations for 30 minutes with vigorous shaking. Primary antibodies were detected using horseradish peroxidase-conjugated species-specific secondary antibodies (1:5000, Cell Signaling Technology) and Clarity ECL blotting substrate (Bio-Rad) or Clarity Max ECL blotting substrate (Bio-Rad). Imaging of blots were performed using the ChemiDoc XRS+ imaging system (Bio-Rad), and quantitative densitometry was performed using the Bio-Rad ImageLab software.

---

**Extended Data Figure 1.**
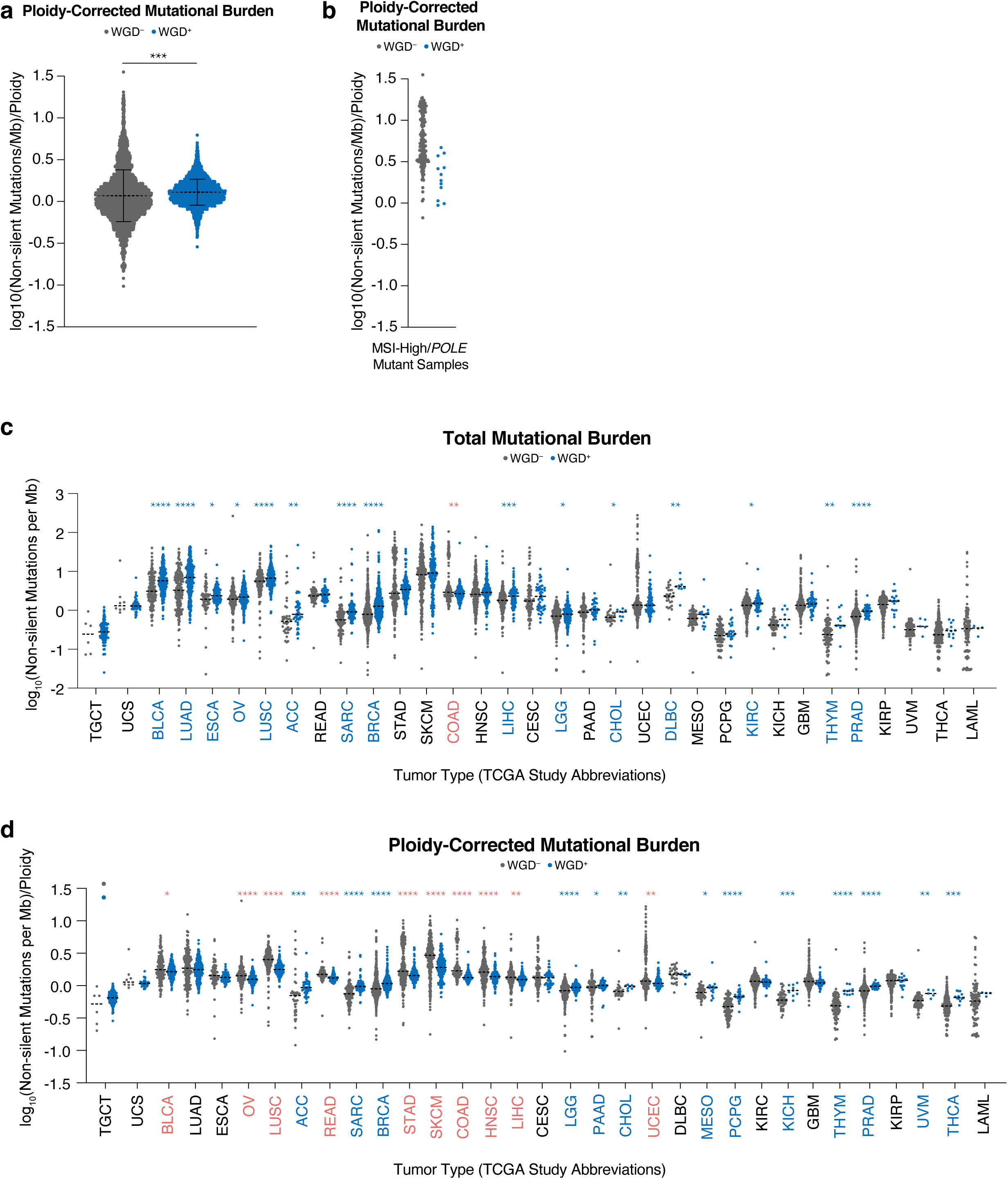
**(a)** Ploidy-corrected mutational burden in WGD^+^ and WGD^−^ samples in the TCGA (multi-variable linear regression; dotted line shows mean +/- SD). **(b)** Ploidy-corrected mutational burden of WGD^+^ and WGD^−^ samples in the TCGA with MSI/*POLE* mutations. **(c)** Total mutational burden in indicated TCGA samples (dotted lines show median; Wilcoxon rank-sum test: green stars indicate higher burden in WGD^−^ samples and blue stars indicate higher burden in WGD^+^ samples). **(d)** Ploidy-corrected mutational burden in indicated TCGA samples (dotted lines show median; Wilcoxon rank-sum test: red stars indicate higher burden in WGD^−^ samples and blue stars indicate higher burden in WGD^+^ samples). * p < 0.05, ** p < 0.01, *** p < 0.001, **** p < 0.0001

**Extended Data Figure 2.**
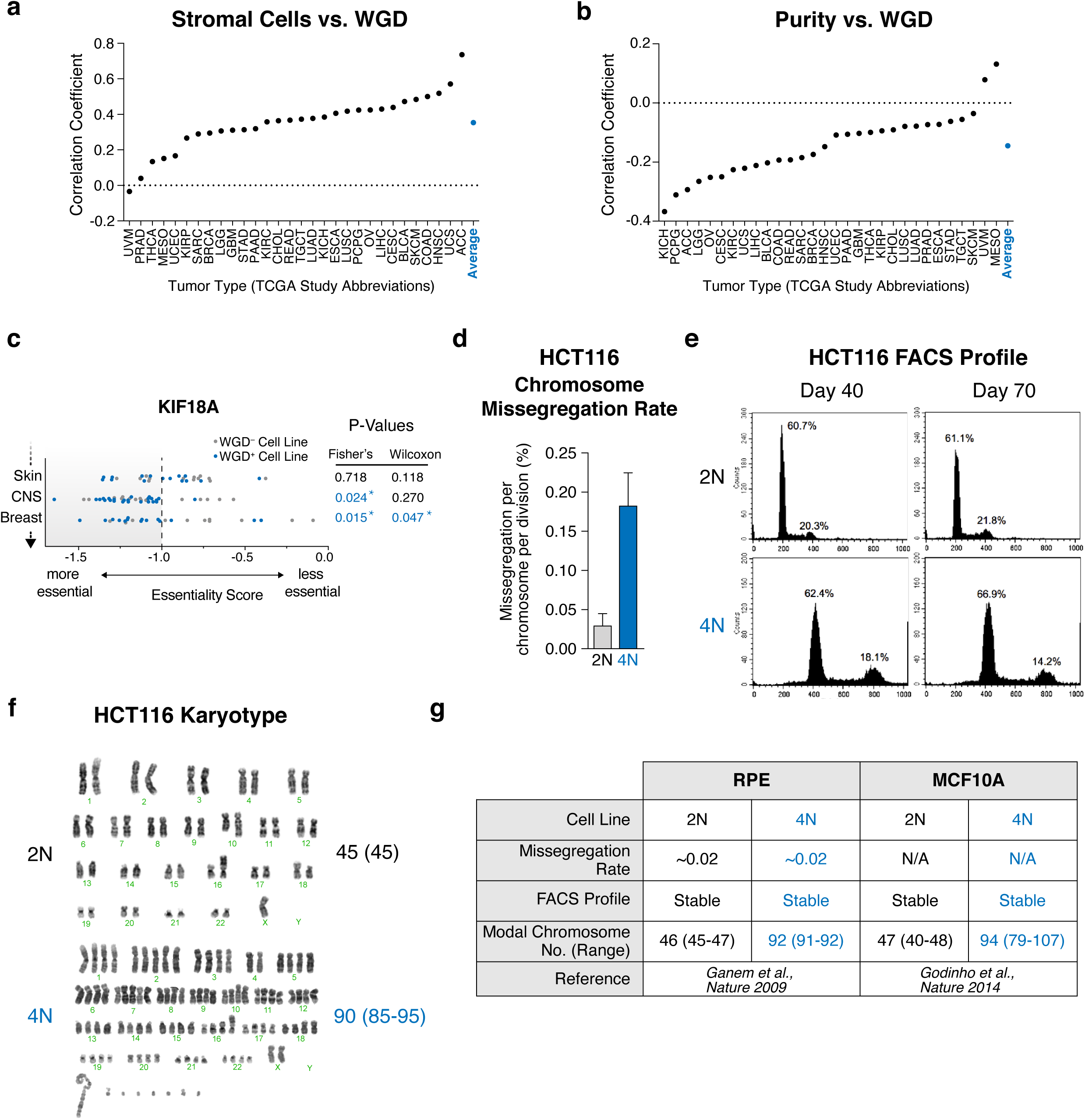
**(a)** Correlation of stromal cell fraction and WGD (Pearson’s correlation). **(b)** Correlation of purity and WGD (Pearson’s correlation). **(c)** Illustration of our ploidy-specific lethal (PSL) analysis using gene essentiality scores for KIF18A across cell lines in three tumor types in the Project Achilles CRISPR dataset. Starred p-values in blue represent instances where the cutoff for enrichment in WGD^+^ cell lines was met in either our thresholded (Fisher’s exact) or non-thresholded (Wilcoxon) analyses (*see methods*). **(d)** HCT116 chromosome missegregation rate (graph shows mean +/- SD). **(e)** DNA FACS profile of diploid and tetraploid HCT116 cells at 40 and 70 days of culture. **(f)** Karyotype of diploid and tetraploid HCT-116 cells with modal chromosome number and range (n = 20 karyotypes analyzed per condition). **(g)** Previously published data demonstrating the stability of isogenic diploid and tetraploid RPE and MCF10A cell lines.

**Extended Data Figure 3.**
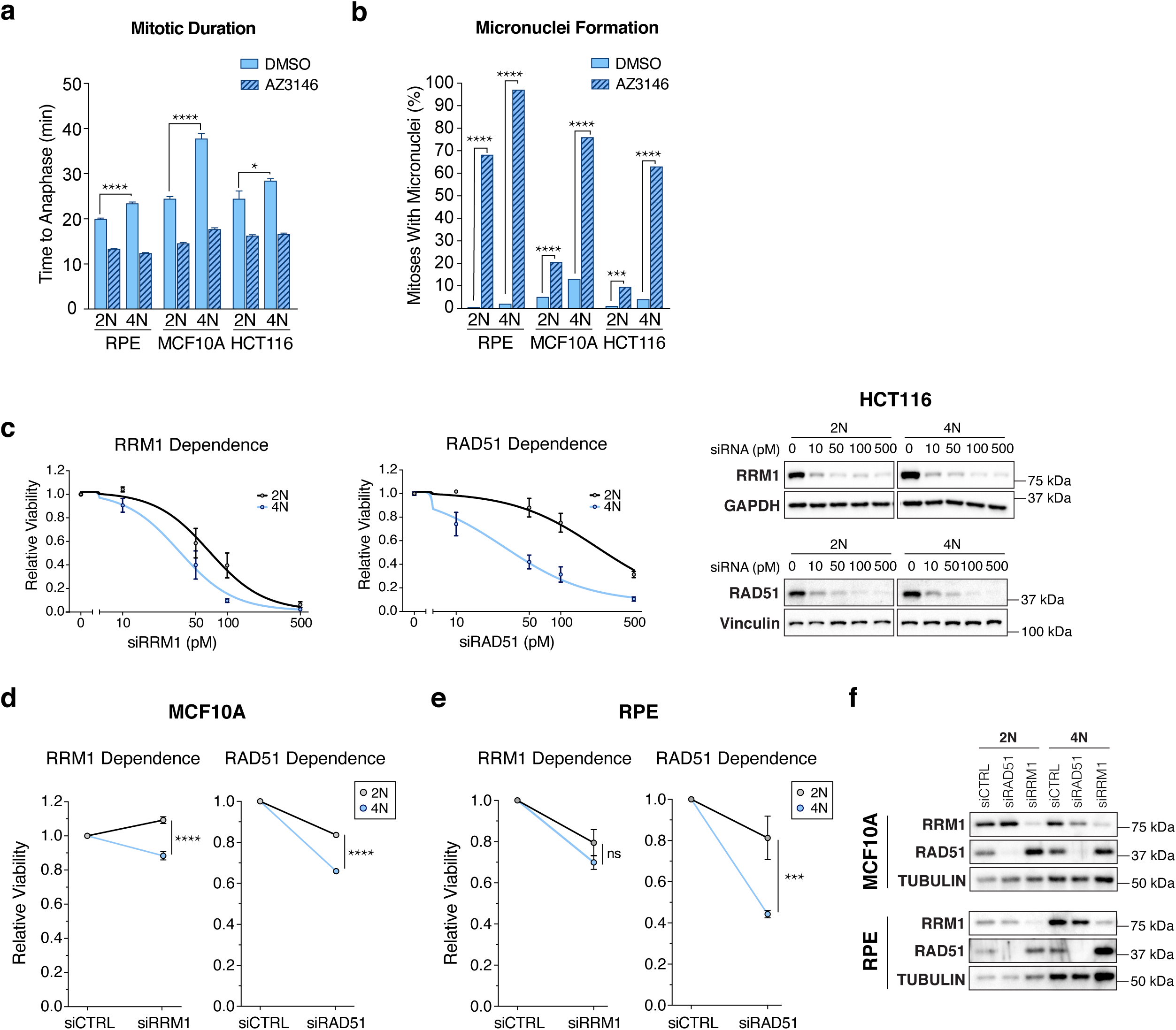
**(a)** Mitotic duration of indicated cells following indicated treatments (n = 200 cells; Student’s unpaired t-test; graph shows mean +/- SEM). **(b)** The fraction of mitoses that generate micronuclei following indicated treatments (n = 200 cells; Student’s unpaired t-test). **(c)** Relative viability of 2N and 4N HCT116 cells 7 days after treatment with indicated siRNA at indicated concentrations with Western blot showing protein knockdown 48 hours after treatment with siRNA (graph shows mean +/- SEM at each dose). **(d)** Relative viability of 2N and 4N MCF10A cells 7 days after treatment with indicated siRNA at 50 pM concentration (Student’s unpaired one-tailed t-test; graph shows mean +/- SEM). **(e)** Relative viability of 2N and 4N RPE cells 5 days after treatment with indicated siRNA at 50 pM concentration (Student’s unpaired one-tailed t-test; graph shows mean +/- SEM). **(f)** Western blot showing knockdown of indicated proteins 48 hours after treatment with indicated siRNA.

**Extended Data Figure 4.**
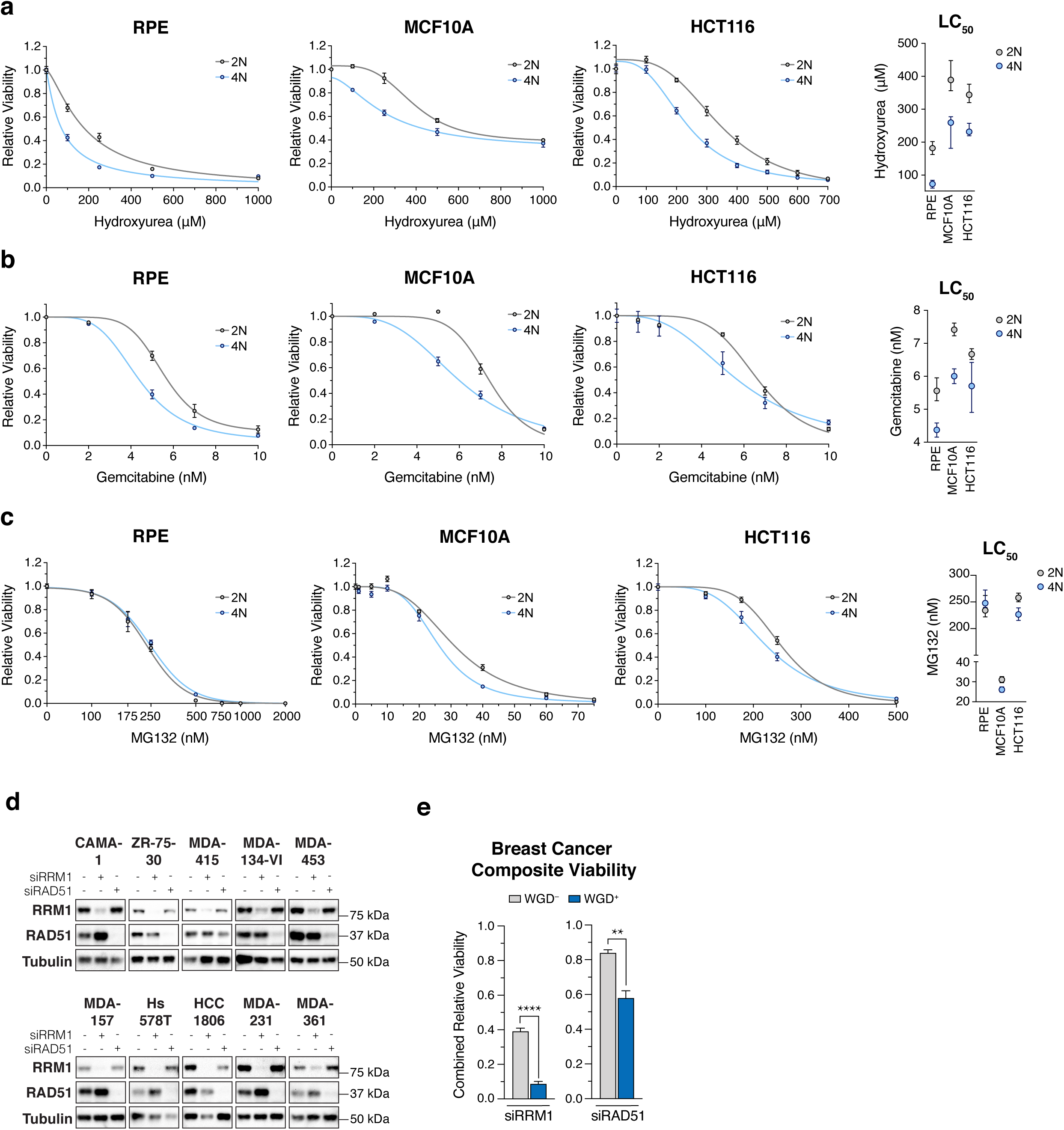
**(a-c)** Dose-response to indicated treatment after 7 days in indicated cell lines with accompanying LC50 (nonlinear regression with variable slope; graph shows mean +/- 95% CI). **(d)** Western blot showing knockdown of indicated proteins in breast cancer cell lines 48 hours after treatment with indicated siRNA. **(e)** Composite viability score of WGD^+^ and WGD^−^ breast cancer cell lines 7 days after treatment with indicated siRNA (Student’s unpaired t-test; graph shows mean +/- SEM). * p < 0.05, ** p < 0.01, *** p < 0.001, **** p < 0.0001:0 01:27 01:48 02:15 02:33

**Extended Data Figure 5.**
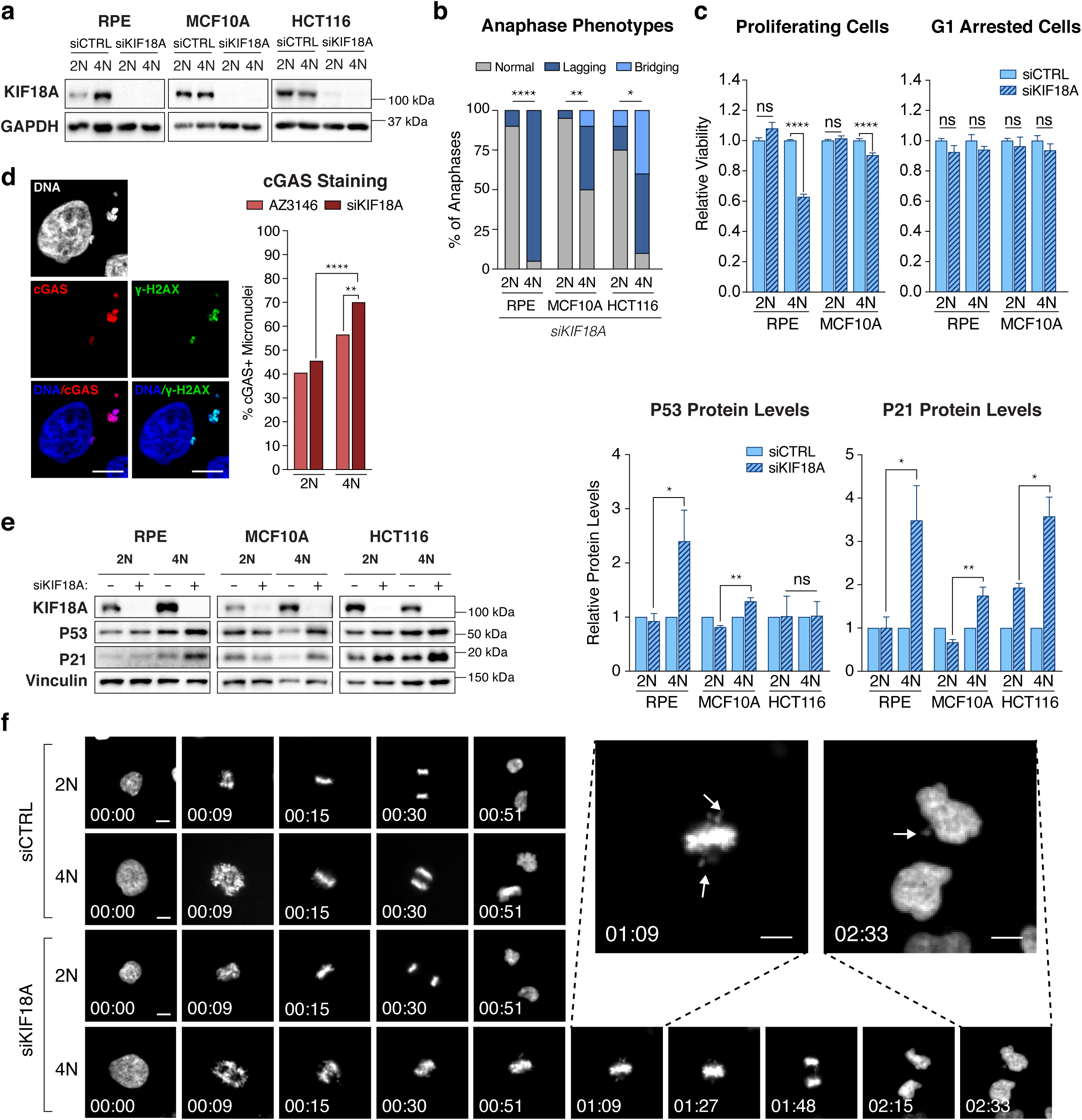
**(a)** Western blot showing KIF18A levels following transfection with the indicated siRNAs in the indicated cell lines. **(b)** Anaphase phenotypes following depletion of KIF18A (n = 20 cells per condition; stars indicate p-value for Fisher’s exact test comparing the fraction of anaphases with lagging chromosomes). **(c)** Relative viability of indicated cell lines 4 days after transfection with the indicated siRNA (Student’s unpaired t-test; graph shows mean +/- SEM). **(d)** Representative image of a 4N MCF10A cell 4 days after transfection with siKIF18A and stained for cGAS. Graph shows the fraction of micronuclei in 2N and 4N MCF10A cells with indicated treatment that stained positive for cGAS (n = 200 micronuclei per condition; Fisher’s exact test; scale bar 10 μm). **(e)** Representative Western blot of indicated protein levels after treatment with indicated siRNA and accompanying graphs showing relative protein levels normalized to loading control (Student’s unpaired one-tailed t-test; graph shows means +/- SEM). **(f)** Representative still images from 2N and 4N MCF10A cells progressing through mitosis after transfection with the indicated siRNAs. H2B-GFP labeled chromosomes are shown in white. Arrows in enlarged images show oscillating chromosomes during metaphase and the generation of a micronucleus (hrs: min; scale bar 10 μm). * p < 0.05, ** p < 0.01, *** p < 0.001, **** p < 0.0001

**Extended Data Figure 6.**
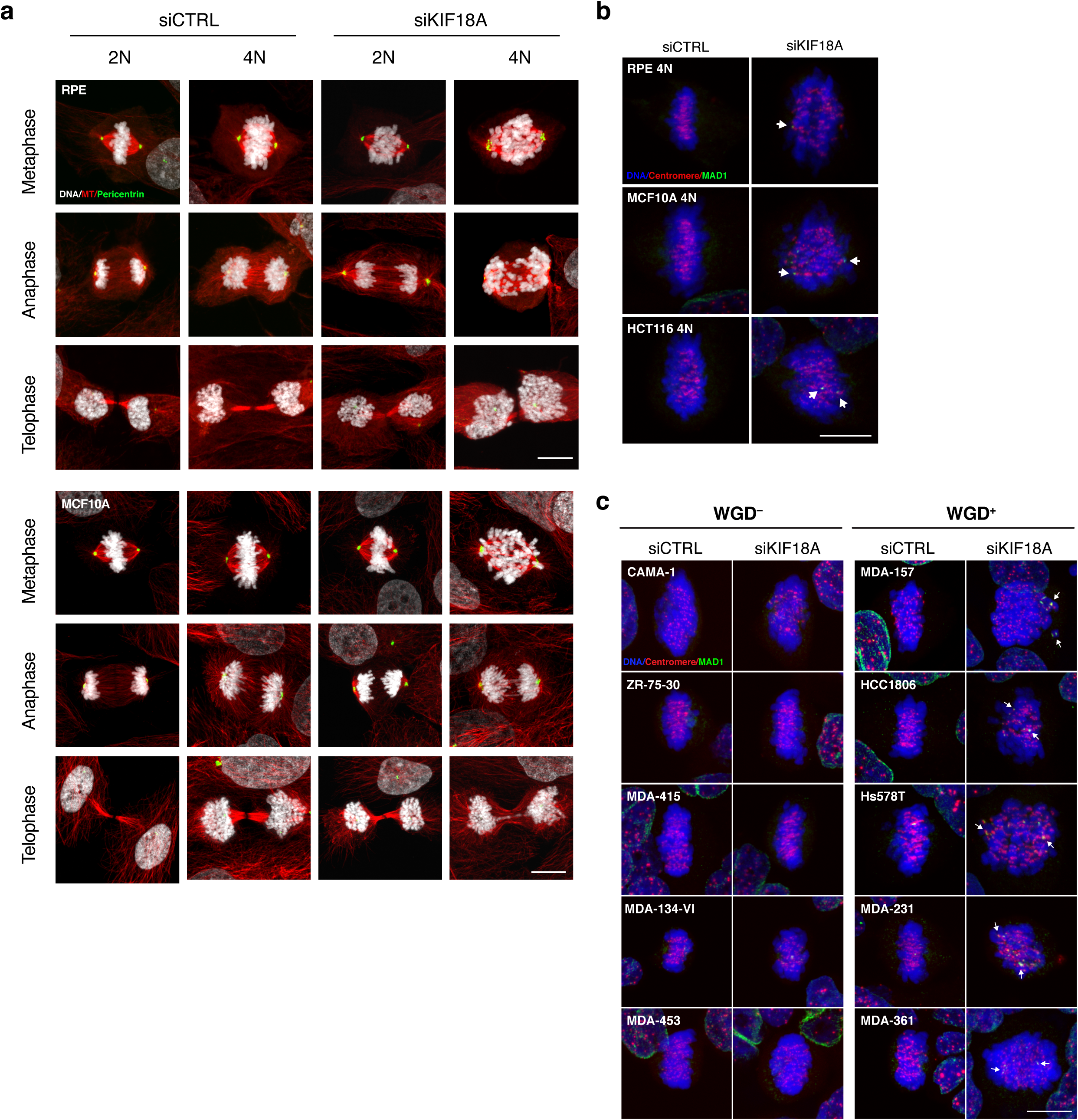
**(a)** Representative confocal images showing phases of mitosis in indicated cell lines 48 hours after transfection with indicated siRNA (scale bar 10 μm). **(b-c)** Representative confocal images pf indicated cell lines 48 hours after transfection with indicated siRNA. Arrows highlight MAD1 positive kinetochores in misaligned chromosomes (scale bar 10μm).

**Extended Data Figure 7.**
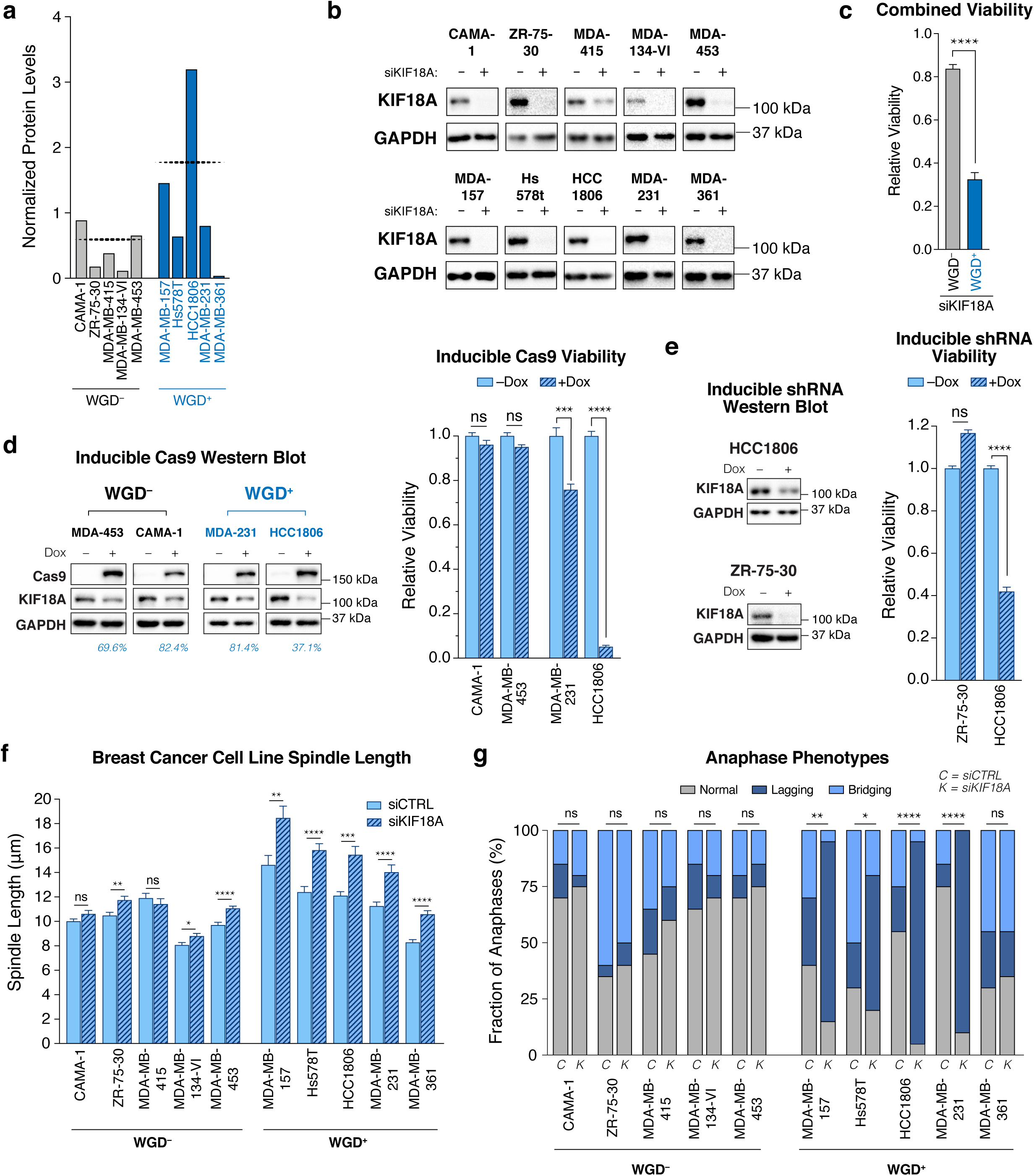
**(a)** Normalized KIF18A protein levels in indicated cell lines (dotted line represents mean). **(b)** Western blot showing KIF18A levels 48 hours after transfection with indicated siRNA. **(c)** Average viability of WGD^+^ and WGD^−^ breast cancer cell lines 7 days after transfection with indicated siRNA (Student’s unpaired t-test). **(d)** Relative viability 7 days after induction of Cas9 in cells with sgRNA targeting KIF18A with Western blot showing protein depletion 72 hours after induction (blue numbers represent the percent of protein remaining relative to controls; graph shows mean +/- SEM; Student’s unpaired t-test)**. (e)** Relative viability 7 days after induction of shRNA targeting KIF18A with Western blot showing protein depletion 120 hours after induction (graph shows mean +/- SEM; Student’s upaired one-tailed t-test). **(f)** Measurement of spindle length (centrosome-to-centrosome) after transfection with indicated siRNA (n = 20 cells per condition; Student’s unpaired t-test; graph shows mean +/- SEM). **(g)** Anapahse phenotypes following depletion of KIF18A (n = 20 cells per condition; stars indicate p-value for Fisher’s exact test comparing the fraction of anaphases with lagging chromosomes). * p < 0.05, ** p < 0.01, *** p < 0.001, **** p < 0.0001

## Methods References

1 Knijnenburg, T. A. et al. Genomic and Molecular Landscape of DNA Damage Repair Deficiency across The Cancer Genome Atlas. Cell Reports 23, 239-254.e236, doi: https://doi.org/10.1016/j.celrep.2018.03.076 (2018).

2 Bailey, M. H. et al. Comprehensive Characterization of Cancer Driver Genes and Mutations. Cell 173, 371-385.e318, doi: 10.1016/j.cell.2018.02.060 (2018).

3 Ellrott, K. et al. Scalable Open Science Approach for Mutation Calling of Tumor Exomes Using Multiple Genomic Pipelines. Cell Systems 6, 271-281.e277, doi: 10.1016/j.cels.2018.03.002 (2018).

4 Carter, S. L. et al. Absolute quantification of somatic DNA alterations in human cancer. Nature Biotechnology 30, 413, doi: 10.1038/nbt.2203 (2015).

5 Taylor, A. M. et al. Genomic and Functional Approaches to Understanding Cancer Aneuploidy. Cancer Cell 33, 676-689.e673, doi: 10.1016/j.ccell.2018.03.007 (2018).

6 Thorsson, V. et al. The Immune Landscape of Cancer. Immunity 48, 812-830.e814, doi: https://doi.org/10.1016/j.immuni.2018.03.023 (2018).

7 McFarland, J. M. et al. Improved estimation of cancer dependencies from large-scale RNAi screens using model-based normalization and data integration. Nature Communications 9, 4610, doi: 10.1038/s41467-018-06916-5 (2018).

8 Meyers, R. M. et al. Computational correction of copy number effect improves specificity of CRISPR–Cas9 essentiality screens in cancer cells. Nature Genetics 49, 1779, doi: 10.1038/ng.3984

9 Fonseca, C. L. et al. Mitotic chromosome alignment ensures mitotic fidelity by promoting interchromosomal compaction during anaphase. The Journal of Cell Biology 218, 1148, doi: 10.1083/jcb.201807228 (2019).

10 Janssen, L. M. E. et al. Loss of Kif18A Results in Spindle Assembly Checkpoint Activation at Microtubule-Attached Kinetochores. Current Biology 28, 2685-2696.e2684, doi: 10.1016/j.cub.2018.06.026 (2018).

## References

1 Lens, S. M. A. & Medema, R. H. Cytokinesis defects and cancer. Nature Reviews Cancer 19, 32–45, doi: 10.1038/s41568-018-0084-6 (2019).

2 Ganem, N. J., Storchova, Z. & Pellman, D. Tetraploidy, aneuploidy and cancer. Current Opinion in Genetics & Development 17, 157–162, doi: https://doi.org/10.1016/j.gde.2007.02.011 (2007).

3 Davoli, T. & Lange, T. d. The Causes and Consequences of Polyploidy in Normal Development and Cancer. Annual Review of Cell and Developmental Biology 27, 585–610, doi: 10.1146/annurev-cellbio-092910-154234 (2011).

4 Davoli, T. & de Lange, T. Telomere-driven tetraploidization occurs in human cells undergoing crisis and promotes transformation of mouse cells. Cancer Cell 21, 765–776, doi: 10.1016/j.ccr.2012.03.044 (2012).

5 Fujiwara, T. et al. Cytokinesis failure generating tetraploids promotes tumorigenesis in p53-null cells. Nature 437, 1043, doi: 10.1038/nature04217 https://www.nature.com/articles/nature04217#supplementary-information (2005).

6 Ganem, N. J., Godinho, S. A. & Pellman, D. A mechanism linking extra centrosomes to chromosomal instability. Nature 460, 278, doi: 10.1038/nature08136 https://www.nature.com/articles/nature08136#supplementary-information (2009).

7 Silkworth, W. T., Nardi, I. K., Scholl, L. M. & Cimini, D. Multipolar Spindle Pole Coalescence Is a Major Source of Kinetochore Mis-Attachment and Chromosome Mis-Segregation in Cancer Cells. PLOS ONE 4, e6564, doi: 10.1371/journal.pone.0006564 (2009).

8 Thompson, D. A., Desai, M. M. & Murray, A. W. Ploidy Controls the Success of Mutators and Nature of Mutations during Budding Yeast Evolution. Current Biology 16, 1581–1590, doi: https://doi.org/10.1016/j.cub.2006.06.070 (2006).

9 Dewhurst, S. M. et al. Tolerance of Whole-Genome Doubling Propagates Chromosomal Instability and Accelerates Cancer Genome Evolution. Cancer Discovery 4, 175, doi: 10.1158/2159-8290.CD-13-0285 (2014).

10 López, S. et al. Interplay between whole-genome doubling and the accumulation of deleterious alterations in cancer evolution. Nature Genetics 52, 283–293, doi: 10.1038/s41588-020-0584-7 (2020).

11 Selmecki, A. M. et al. Polyploidy can drive rapid adaptation in yeast. Nature 519, 349–352, doi: 10.1038/nature14187 (2015).

12 Bielski, C. M. et al. Genome doubling shapes the evolution and prognosis of advanced cancers. Nature Genetics, doi: 10.1038/s41588-018-0165-1 (2018).

13 Priestley, P. et al. Pan-cancer whole-genome analyses of metastatic solid tumours. Nature 575, 210–216, doi: 10.1038/s41586-019-1689-y (2019).

14 Ganem, Neil J. et al. Cytokinesis Failure Triggers Hippo Tumor Suppressor Pathway Activation. Cell 158, 833–848, doi: https://doi.org/10.1016/j.cell.2014.06.029 (2014).

15 Senovilla, L. et al. An Immunosurveillance Mechanism Controls Cancer Cell Ploidy. Science 337, 1678, doi: 10.1126/science.1224922 (2012).

16 Andreassen, P. R., Lohez, O. D., Lacroix, F. B. & Margolis, R. L. Tetraploid State Induces p53-dependent Arrest of Nontransformed Mammalian Cells in G1. Molecular Biology of the Cell 12, 1315–1328, doi: 10.1091/mbc.12.5.1315 (2001).

17 Ben-David, U. & Amon, A. Context is everything: aneuploidy in cancer. Nature Reviews Genetics, doi: 10.1038/s41576-019-0171-x (2019).

18 Potapova, T. A., Seidel, C. W., Box, A. C., Rancati, G. & Li, R. Transcriptome analysis of tetraploid cells identifies cyclin D2 as a facilitator of adaptation to genome doubling in the presence of p53. Molecular Biology of the Cell 27, 3065–3084, doi: 10.1091/mbc.e16-05-0268 (2016).

19 Crockford, A. et al. Cyclin D mediates tolerance of genome-doubling in cancers with functional p53. Annals of Oncology 28, 149–156, doi: 10.1093/annonc/mdw612 (2016).

20 Storchová, Z. et al. Genome-wide genetic analysis of polyploidy in yeast. Nature 443, 541–547, doi: 10.1038/nature05178 (2006).

21 Lin, H. et al. Polyploids require Bik1 for kinetochore-microtubule attachment. The Journal of cell biology 155, 1173–1184, doi: 10.1083/jcb.200108119 (2001).

22 Coward, J. & Harding, A. Size Does Matter: Why Polyploid Tumor Cells are Critical Drug Targets in the War on Cancer. Frontiers in Oncology 4, doi: 10.3389/fonc.2014.00123 (2014).

23 Carter, S. L. et al. Absolute quantification of somatic DNA alterations in human cancer. Nature Biotechnology 30, 413, doi: 10.1038/nbt.2203 https://www.nature.com/articles/nbt.2203#supplementary-information (2012).

24 Zack, T. I. et al. Pan-cancer patterns of somatic copy number alteration. Nature Genetics 45, 1134, doi: 10.1038/ng.2760 https://www.nature.com/articles/ng.2760#supplementary-information (2013).

25 Shlien, A. et al. Combined hereditary and somatic mutations of replication error repair genes result in rapid onset of ultra-hypermutated cancers. Nature Genetics 47, 257–262, doi: 10.1038/ng.3202 (2015).

26 Ciriello, G. et al. Emerging landscape of oncogenic signatures across human cancers. Nature Genetics 45, 1127–1133, doi: 10.1038/ng.2762 (2013).

27 Chalmers, Z. R. et al. Analysis of 100,000 human cancer genomes reveals the landscape of tumor mutational burden. Genome Medicine 9, 34, doi: 10.1186/s13073-017-0424-2 (2017).

28 Antao, N. V., Marcet-Ortega, M., Cifani, P., Kentsis, A. & Foley, E. A. A Cancer-Associated Missense Mutation in PP2A-Aα Increases Centrosome Clustering during Mitosis. iScience 19, 74–82, doi: https://doi.org/10.1016/j.isci.2019.07.018 (2019).

29 Thorsson, V. et al. The Immune Landscape of Cancer. Immunity 48, 812-830.e814, doi: https://doi.org/10.1016/j.immuni.2018.03.023 (2018).

30 Aran, D., Hu, Z. & Butte, A. J. xCell: digitally portraying the tissue cellular heterogeneity landscape. Genome Biology 18, 220, doi: 10.1186/s13059-017-1349-1 (2017).

31 Taylor, A. M. et al. Genomic and Functional Approaches to Understanding Cancer Aneuploidy. Cancer Cell 33, 676-689.e673, doi: 10.1016/j.ccell.2018.03.007 (2018).

32 Davoli, T., Uno, H., Wooten, E. C. & Elledge, S. J. Tumor aneuploidy correlates with markers of immune evasion and with reduced response to immunotherapy. Science 355, doi: 10.1126/science.aaf8399 (2017).

33 McFarland, J. M. et al. Improved estimation of cancer dependencies from large-scale RNAi screens using model-based normalization and data integration. Nature Communications 9, 4610, doi: 10.1038/s41467-018-06916-5 (2018).

34 Meyers, R. M. et al. Computational correction of copy number effect improves specificity of CRISPR–Cas9 essentiality screens in cancer cells. Nature Genetics 49, 1779, doi: 10.1038/ng.3984 https://www.nature.com/articles/ng.3984#supplementary-information (2017).

35 Broad, D. DepMap Achilles 18Q3 Public. (2018).

36 Godinho, S. A. et al. Oncogene-like induction of cellular invasion from centrosome amplification. Nature 510, 167–171, doi: 10.1038/nature13277 (2014).

37 Musacchio, A. The Molecular Biology of Spindle Assembly Checkpoint Signaling Dynamics. Current Biology 25, R1002–R1018, doi: https://doi.org/10.1016/j.cub.2015.08.051 (2015).

38 Yang, Z., Lončarek, J., Khodjakov, A. & Rieder, C. L. Extra centrosomes and/or chromosomes prolong mitosis in human cells. Nature Cell Biology 10, 748–751, doi: 10.1038/ncb1738 (2008).

39 Mohamed Jemaa, G. M., Gwendaline Lledo, Delphine Lissa, Christelle Reynes, Nathalie Morin, Frederic Chibon, Antonella Sistigu, Maria Castedo, Ilio Vitale, Guido Kroemer, Ariane Abrieu. Whole-genome duplication increases tumor cell sensitivity to MPS1 inhibition. Oncotarget, 885–901 (2016).

40 Vitale, I. et al. Inhibition of Chk1 Kills Tetraploid Tumor Cells through a p53-Dependent Pathway. PLOS ONE 2, e1337, doi: 10.1371/journal.pone.0001337 (2007).

41 Wangsa, D. et al. Near-tetraploid cancer cells show chromosome instability triggered by replication stress and exhibit enhanced invasiveness. The FASEB Journal 32, 3502–3517, doi: 10.1096/fj.201700247RR (2018).

42 Zheng, L. et al. Polyploid cells rewire DNA damage response networks to overcome replication stress-induced barriers for tumour progression. Nature Communications 3, 815, doi: 10.1038/ncomms1825 (2012).

43 Jordheim, L. P., Sève, P., Trédan, O. & Dumontet, C. The ribonucleotide reductase large subunit (RRM1) as a predictive factor in patients with cancer. The Lancet Oncology 12, 693–702, doi: https://doi.org/10.1016/S1470-2045(10)70244-8 (2011).

44 Santaguida, S. & Amon, A. Short- and long-term effects of chromosome mis-segregation and aneuploidy. Nature Reviews Molecular Cell Biology 16, 473–485, doi: 10.1038/nrm4025 (2015).

45 Stumpff, J., von Dassow, G., Wagenbach, M., Asbury, C. & Wordeman, L. The Kinesin-8 Motor Kif18A Suppresses Kinetochore Movements to Control Mitotic Chromosome Alignment. Developmental Cell 14, 252–262, doi: https://doi.org/10.1016/j.devcel.2007.11.014 (2008).

46 Stumpff, J., Wagenbach, M., Franck, A., Asbury, C. L. & Wordeman, L. Kif18A and chromokinesins confine centromere movements via microtubule growth suppression and spatial control of kinetochore tension. Developmental cell 22, 1017–1029, doi: 10.1016/j.devcel.2012.02.013 (2012).

47 Fonseca, C. L. et al. Mitotic chromosome alignment ensures mitotic fidelity by promoting interchromosomal compaction during anaphase. The Journal of Cell Biology 218, 1148, doi: 10.1083/jcb.201807228 (2019).

48 Mayr, M. I. et al. The Human Kinesin Kif18A Is a Motile Microtubule Depolymerase Essential for Chromosome Congression. Current Biology 17, 488–498, doi: 10.1016/j.cub.2007.02.036 (2007).

49 Czechanski, A. et al. Kif18a is specifically required for mitotic progression during germ line development. Developmental Biology 402, 253–262, doi: https://doi.org/10.1016/j.ydbio.2015.03.011 (2015).

50 Reinholdt, L. G., Munroe, R. J., Kamdar, S. & Schimenti, J. C. The mouse gcd2 mutation causes primordial germ cell depletion. Mechanisms of Development 123, 559–569, doi: https://doi.org/10.1016/j.mod.2006.05.003 (2006).

51 Hatch, Emily M., Fischer, Andrew H., Deerinck, Thomas J. & Hetzer, Martin W. Catastrophic Nuclear Envelope Collapse in Cancer Cell Micronuclei. Cell 154, 47–60, doi: 10.1016/j.cell.2013.06.007 (2013).

52 Mackenzie, K. J. et al. cGAS surveillance of micronuclei links genome instability to innate immunity. Nature 548, 461–465, doi: 10.1038/nature23449 (2017).

53 Harding, S. M. et al. Mitotic progression following DNA damage enables pattern recognition within micronuclei. Nature 548, 466–470, doi: 10.1038/nature23470 (2017).

54 Zhang, C.-Z. et al. Chromothripsis from DNA damage in micronuclei. Nature 522, 179, doi: 10.1038/nature14493 https://www.nature.com/articles/nature14493#supplementary-information (2015).

55 Janssen, L. M. E. et al. Loss of Kif18A Results in Spindle Assembly Checkpoint Activation at Microtubule-Attached Kinetochores. Current Biology 28, 2685-2696.e2684, doi: 10.1016/j.cub.2018.06.026 (2018).

56 Edzuka, T. & Goshima, G. Drosophila kinesin-8 stabilizes the kinetochore–microtubule interaction. The Journal of Cell Biology 218, 474–488, doi: 10.1083/jcb.201807077 (2018).

57 Zhu, H. et al. Targeted deletion of Kif18a protects from colitis-associated colorectal (CAC) tumors in mice through impairing Akt phosphorylation. Biochemical and Biophysical Research Communications 438, 97–102, doi: https://doi.org/10.1016/j.bbrc.2013.07.032 (2013).

58 Zhang, C. et al. Kif18A is involved in human breast carcinogenesis. Carcinogenesis 31, 1676–1684, doi: 10.1093/carcin/bgq134 (2010).

